# Perturbation analysis of a multi-morphogen Turing Reaction-Diffusion stripe patterning system reveals key regulatory interactions

**DOI:** 10.1101/2019.12.27.889493

**Authors:** Andrew D. Economou, Nicholas A.M. Monk, Jeremy B.A. Green

## Abstract

Periodic patterning is extremely widespread in developmental biology and is broadly modelled by Reaction-Diffusion (RD) processes. However, the minimal two-component RD system is vastly simpler than the multimolecular events that current biology is now able to describe. Moreover, RD models are typically underconstrained such that it is often hard to meaningfully relate the model architecture to real interactions measured experimentally. To address both these issues, we investigated the periodic striped patterning of the rugae (transverse ridges) in the roof of the mammalian palate. We experimentally implicated a small number of major signalling pathways and established theoretical limits on the number of pathway network topologies that can account for the stable spatial phase relationships of their observed signalling outputs. We further conducted perturbation analysis both experimentally and in silico, critically to assess the effects of perturbations on established patterns. We arrived at a relatively highly-constrained number of possible networks and found that these share some common motifs. Finally, we examined the dynamics of pattern appearance and discovered a core network consisting of epithelium-specific FGF and Wnt as mutually antagonistic “activators” and Shh as the “inhibitor”, which initiates the periodicity and whose existence constrains the network topology still further. Together these studies articulate the principles of multi-morphogen RD patterning and demonstrate the utility of perturbation analysis as a tool for constraining networks in this and, in principle, any RD system.

## Introduction

The generation of anatomy by self-organisation remains one of the most important subjects in the study of biology. It has acquired new importance as a guiding feature of regenerative medicine and the modelling of disease processes by the creation of self-organising organoids from stem cells (Werner, Vu, & Rink, 2017). Setting aside mechanical self-organisation, a central idea of how self-organisation occurs chemically is that of Reaction-Diffusion (RD). In Turing’s initial formulation, an initially stable system of two or more interacting morphogens can be destabilised (i.e. generate non-uniform distributions) into a periodic pattern through diffusion (Turing, 1952). This is referred to as a diffusion-driven instability (DDI). Gierer and Meinhardt later refined Turing’s ideas and, among several refinements, introduced the formulation of the RD system as a system consisting of a slow-diffusing activator and fast-diffusing inhibitor (Meinhardt, 2012). Since the 1970s, numerous examples of RD behaviour have been described and analysed, and more recently the methods of molecular biology and biochemistry have identified a number of morphogen pairs – generally protein growth factors or growth-factor-binding proteins – that fit Turing’s and Meinhardt’s minimal description (e.g. Economou et al., 2012; Jung et al., 1998; Michon, Forest, Collomb, Demongeot, & Dhouailly, 2008; Mou, Jackson, Schneider, Overbeek, & Headon, 2006; Sick, Reinker, Timmer, & Schlake, 2006).

One aspect of self-organisation that has been set aside hitherto is that, as we now know, large proportions of the genome and proteome are devoted to regulation. Consequently, minimal systems must be expanded if they are to capture this complexity where it is functionally relevant. The extension of RD systems to include multiple morphogens and even non-diffusible components (Celliere, Menshykau, & Iber, 2012; Klika, Baker, Headon, & Gaffney, 2012; Marcon, Diego, Sharpe, & Muller, 2016; Raspopovic, Marcon, Russo, & Sharpe, 2014) opens up a Pandora’s box of possible descriptions of self-patterning systems. It raises the question of what level of description is usefully interpretable and, importantly, experimentally tractable, such that empirical data and theory can be compared. It may be argued that conceptually and pharmacologically, the most intelligible and accessible level of description is the signalling pathway: typically, this puts a family of protein growth factor ligands (e.g. the FGFs) together with their common receptors, transducers and target genes into a single unit. This provides a potentially happy balance between the minimalism of a pure two-component morphogen system and the overwhelming complexity of an ‘omic molecular network.

Taking this level as our motivating principle, we here ask how far one can get in defining a model – a network topology – of key interactions that can capture the behaviour of the system. We analyse the periodic pattern that generates the transverse ridges on the roof of the mouse palate, the rugae, a striped pattern that we previously showed to exhibit RD system behaviour (Economou et al., 2012). Using chemical inhibitors and pathway target expression analysis together with analytical and numerical simulation strategies, we have constrained the model possibilities to a small number of consistent topologies. In doing so, we identify a useful theory-experiment dialogue that generates specific hypotheses amenable to practical progress in understanding the dynamic behaviours of this example of a self-organising RD system.

## Materials & Methods

### Generation of embryos & explants

Wild type CD1 embryos were harvested, staged, fixed and stained for wholemount *in situ* hybridization using established methods. Palate explants from embryonic day 13.5 (E13.5) embryos were made using 0.1 mm tungsten needles and cultured at 37°C in 5% CO_2_ atmosphere for 24 hours using the Trowell technique (Alfaqeeh & Tucker, 2013) in serum-free Advanced D-MEM/F12 (GibcoBRL), 20 U/ml Pen-strep (GibcoBRL), 50 mM transferrin (Sigma) and 150 μg/ml ascorbic acid (Sigma). Chemical inhibitors were added at the beginning of the 24 hour culture period at following final concentrations: SU5402 (Calbiochem) at 40 μM; Cyclopamine (Sigma) at 20 μM; IWP-2 (Cambridge Bioscience/Cell Guidance Systems) at 50 μM; Dorsomorphin (Cayman Chemical) at 50 μM. Control palates were contralateral explants from the same embryo incubated with vehicle only. Experiments were repeated at least four times for each condition.

### In situ hybridization

For section in situ hybridization, fixed specimens were embedded in wax and serially sectioned at 7 μm thickness, with successive sections mounted on four different slides to allow different probes to be used on nearby/adjacent sections. Wholemount and section in situ hybridisation was conducted according to standard methods. Probes were gifts of colleagues obtained ultimately from authors of published references as follows: *Shh* (Echelard et al., 1993); *Lef1* (Gat, DasGupta, Degenstein, & Fuchs, 1998); *Id1* (Rice et al., 2000); *Spry2*(Tefft et al., 1999); *Ptc1* (Goodrich, Jung, Higgins, & Scott, 1999); *Gli1* (Hui, Slusarski, Platt, Holmgren, & Joyner, 1994); *Etv4* (*Pea3*) *and Etv5* (*Erm1*) (Chotteau-Lelievre, Desbiens, Pelczar, Defossez, & de Launoit, 1997); *Axin2*(Lustig et al., 2002). Whole stained explants placed in a minimum volume of PBS in wells cut into 1% agarose were digitally imaged under a stereo dissecting microscope.

### Image analysis

The mouse embryonic head was sectioned at 7 μm intervals in the sagittal aspect. For a given marker, in situ hybridization was performed on every fourth or fifth section, for the entire mediolateral extent of the rugae. Sections were imaged using a Zeiss Axioskop upright microscope using a 20X objective under both brightfield and phase contrast. To record variation of relative expression along the anteroposterior axis of the palatal epithelium and underlying mesenchyme, the apical and basal extents of the epithelium were first manually traced on the phase contrast images (where unstained epithelium could be clearly seen) using ImageJ. We wrote a macro in ImageJ (see link in Supplementary Information) to return the relative staining intensity (as recorded in the brightfield images converted to 8-bit greyscale) within the traces along the epithelium (averaged across its thickness) and in the underlying 25 μm of mesenchyme. The basal-to-apical thickness of the epithelium was also captured. The position of ruga 3 within the trace was recorded manually. For a given palatal shelf, intensity profiles were made for all sections stained for a particular marker (sections damaged during the sectioning and staining process were excluded). Because the rugae are approximately parallel, it was sufficient to align the intensity profiles to the position of ruga 3 to obtain an average intensity profile along the AP axis of a palatal shelf (Fig.1 supplement 1).

### Construction of kymographs

As the palatal epithelium extends along its AP axis through localized growth immediately anterior to ruga 8, the AP palatal distance between ruga 3 and ruga 8 was used to measure embryonic age (i.e. time). First a high-resolution calibration curve was constructed relating embryonic weights to time-of-harvest for 412 embryos across 38 litters, as in (Peterka, Lesot, & Peterkova, 2002). Weights and rugae 3-to-8 distances of in situ hybridised embryos could then be related to embryonic age (data not shown). Processing to make the kymographs was carried out in R (R Core Team, 2013) (see link to code in supplementary data) as follows. To produce a smooth kymograph, intensity was normalized (scaled) to values between 0 and 1, positioned by time, and aligned in the AP direction at ruga 3. A moving average (using a 0.2 day window) was calculated to smooth the intensity plot in the time axis. Where there were fewer than 2 traces in the window, no value was plotted. Finally, the kymograph was replotted as distance relative to ruga 8.

### Numerical simulations

Reaction-diffusion systems were simulated in R using a piecewise linear model (code available through link in Supplementary Information) of the form

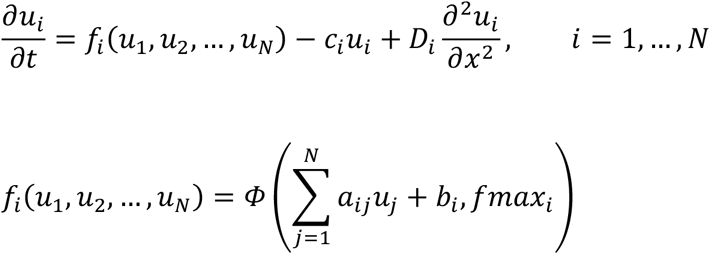

with

*Φ*(*z*, *fmax*) = 0 for *z* < 0
*Φ*(*z*, *fmax*) = *z* for 0 ≤ *z* ≤ *fmax*
*Φ*(*z*, *fmax*) = *fmax* for *z* > *fmax*

The *N* variables *u_i_*(*x, t*), *i* = 1,…,*N*, represent the concentrations of each component of the RD system as functions of time and a single spatial variable *x* (distance along the anteroposterior axis of the palate). The non-negative parameters *c_i_* and *D_i_* represent the degradation rate and diffusion coefficients of the components, respectively.

The function *f_i_*(*u*_1_,…,*u_N_*), *i* = 1,…,*N*, specifies the production rate of component *i*, which takes a non-negative value between 0 and *fmax_i_*. We take this to be a piecewise linear function *Φ* of the weighted sum of all regulatory inputs, where the parameter *a_ij_* represents the weight of the direct regulatory input from component *j* to component *i* (i.e. the sensitivity of the production rate of *i* to changes in the concentration of component *j*.)

The qualitative form (topology) of a RD system is specified by the nature of the interactions between its components (positive, negative, or no interaction). For any given topology, interaction parameters (*a_ij_* where *i* ≠ *j*) were either set to 0 (i.e. no interaction), or assigned a value at random from a uniform distribution between −1 and 1, with the sign determined by the nature (positive or negative) of the interaction. For self-interactions (*a_ij_* where *i* = *j*), the composite parameter *a_ii_*-*c_i_* was drawn from this uniform distribution, taking a negative weighting when there is no autoregulation (*a_ii_*=0). When a component autoactivates, the parameter takes a positive value, with the weight of -*c_i_* also being drawn from the distribution. Diffusion coefficients (*D_i_*) were drawn as the reciprocal of values from a uniform distribution between 1 and 10,000, meaning that at the lower end of the distribution, small differences would have a large effect on the diffusion range.

To assess whether a particular parameterization of the RD system would support the formation of spatial patterns through diffusion driven instability (DDI), and what phase-type of pattern the parameterization could produce, the parameters were compared to the criteria for a DDI generating stable, spatially periodic, non-oscillating patterns described initially by White and Gilligan (White & Gilligan, 1998) with some additional elaboration described in Supplemental Note 1.

To ensure that spatially periodic solutions of the model do not take negative values, they must form around a positive spatially-uniform steady state (all *u_i_* > 0); to ensure this, appropriate production constants *b_i_* are required (Raspopovic et al., 2014). Therefore, for parameter sets that support DDI, constant regulatory input terms (*b_i_*) were calculated so that for each component, the spatially-uniform steady state concentration was fixed at 1. In order for the amplitude of patterns generated from linear RD to remain bounded, the production functions *f_i_*(*u*_1_,…,*u_N_*) must be bounded (Shoji, Iwasa, & Kondo, 2003). We set the lower bound to be zero (production cannot be negative) and the upper bounds to be *fmax_i_*. To ensure that the maximum production rates *fmax_i_* were greater than the production rate at steady state (so that the production rates at steady state are linear functions of their inputs), the *fmax_i_* were set randomly between 1.5 and 3 times the magnitude of *c_i_*.

RD simulations were run using a finite difference scheme with zero flux boundary conditions on a discrete 1D grid of 100 positions. Parameter sets were scaled to ensure non-discretised spatially periodic solutions would be formed on the spatial domain used by comparing the wavelength and growth rate (as detailed in Supplementary Note 1) to values known to fit within the simulation space and time to generate a scaling factor, and then using that factor to run an initial simulation from which spatial patterns could be measured and the parameters scaled more precisely.

### Perturbation analysis

Once a stable spatial pattern was established in the RD system, components were inhibited in one of two different ways. Inhibition of the receptor of a given morphogen (as seen for the inhibitors Cyclopamine, SU5402 and Dorsomorphin, for example) was implemented as an equal proportional reduction of all interaction coefficients representing the response to that morphogens (so if the response to component *u_j_* was inhibited, the values of *a_ij_* were reduced by a factor (1 − *α*), for all values of *i*). Inhibition of the production of a component (as in IWP2, for example) was achieved by proportionately reducing the production term of that component by a factor (1 − *α*) (see **Suppl. Note 2.** for details). For all parameter sets, by successively reducing the strength of the inhibition parameter *α* by a factor of 0.25 from complete inhibition (*α* = 1), we determined empirically for each pathway the maximum perturbation *α_max_* that still allowed patterning but did not perturb the number of whole wavelengths or fail to achieve a stable amplitude. The test statistic for perturbation effects was the mean level for each component relative to an unperturbed run.

### Topology search

For the 3-component RD system, parameter space was systematically sampled to identify parameter sets and topologies capable of giving DDI. Each of the nine reaction parameters *a_ij_* was varied linearly to give a total of 40,353,607 parameter sets. Diffusion parameters were initially set as two fast and one slow or vice versa, although this requirement (based on the longstanding but recently overturned (Marcon et al., 2016) idea that this difference is essential) turned out not to be essential (see Results below). DDI criteria for stable, periodic non-oscillating patterns, as mentioned above and described in detail in Supplemental Note 2 were applied.

## Results

We previously showed that the periodic stripes of expression of the *Sonic hedgehog* (Shh) gene in the mouse mid-gestation palate depend on Shh itself as an “inhibitor” and FGF as an “activator” (Economou et al., 2012). We were deliberately agnostic about the specific FGF ligand-receptor pair that was critical because multiple FGFs and FGF receptors are expressed in the palate (Porntaveetus, Oommen, Sharpe, & Ohazama, 2010). Both FGF and Shh are, by a number of experimental criteria, secreted, diffusible morphogens (Bokel & Brand, 2013; Dessaud et al., 2007). As we acknowledged, there was already published data implicating Wnt and potentially BMP as additional morphogens (Lin et al., 2011; Welsh & O’Brien, 2009). To go beyond the simple two-component description of the patterning network, we sought first to determine the requirement for each of these four morphogen pathways, using an established explant system, as before. We tested the effect of pathway-specific inhibitors on the pattern, specifically using the stripes of Shh expression as our readout (although in principle, any of the components could be used as a readout). Fig.1a shows that inhibition of each of the four pathways has an effect on the *Shh* expression stripes. Dorsomorphin (a BMP pathway inhibitor) intensified and broadened the stripes similar to what we previously reported for cyclopamine (a Hedgehog pathway inhibitor), while IWP-2 (a Wnt pathway inhibitor) narrowed and weakened them. FGF inhibition by SU5402 also weakened the stripes, as previously reported (Economou et al., 2012), but careful inspection over a number of trials also revealed consistent broadening as well weakening (Fig.1A and Fig.1 supplement 2). These results confirm that all four pathways contribute to formation of the ruga pattern.

**Fig 1.**
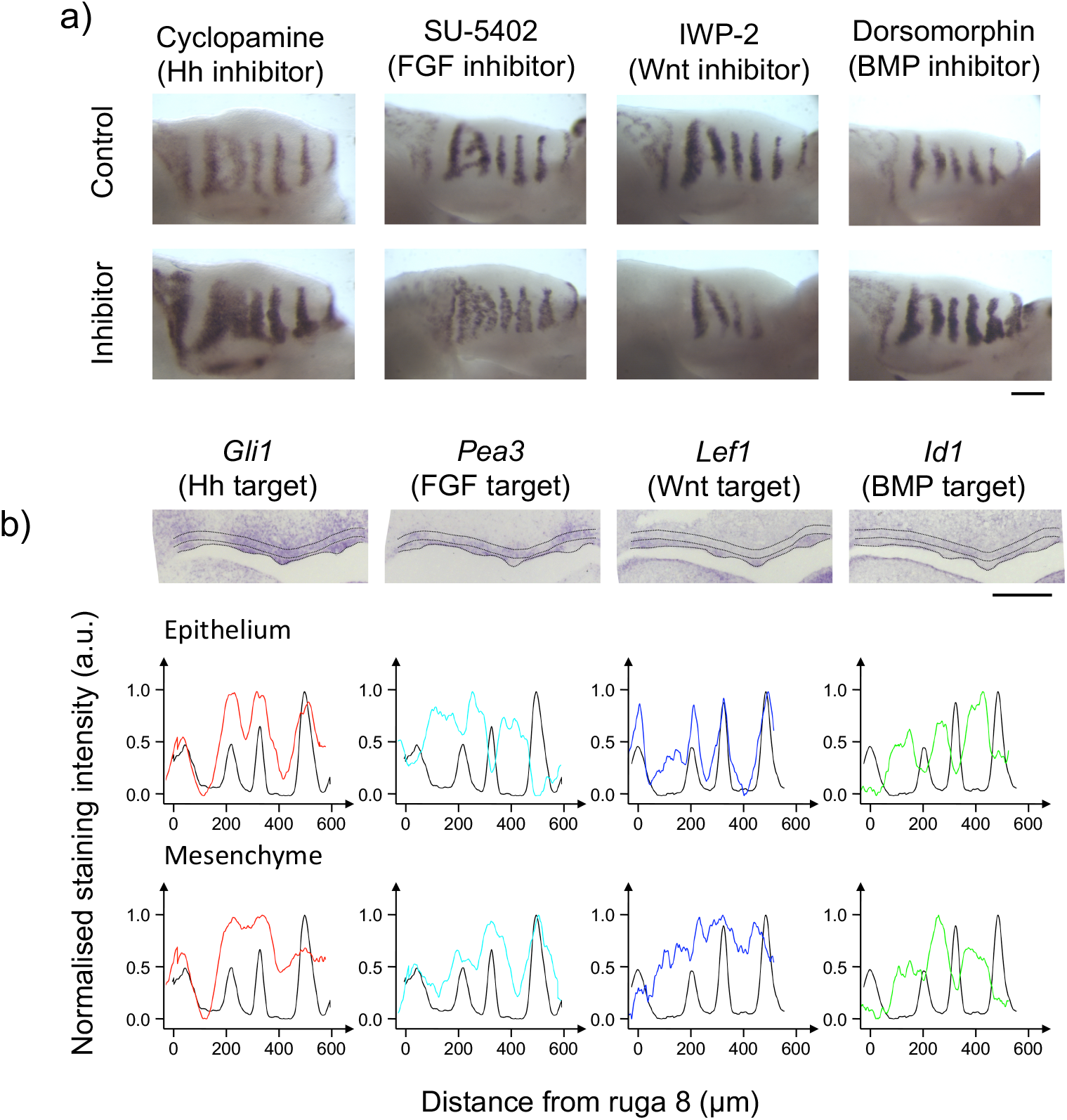
Target expression and responses to inhibitors reveal involvement of Hh, FGF, Wnt and BMP pathways in periodic rugae patterning. a) Shh in situ hybridisations on E13.5 palatal shelf explants cultured for 24 hrs in the specified small molecule inhibitor contralateral shelves as vehicle controls. Anterior to the right, medial up. b) In situ hybridisation of sagittal section through E13.5 palatal shelf for specified genes. Dotted lines illustrate the extent of the palatal epithelium and the underlying mesenchyme used for quantifications. Anterior to the right. The intensity profile averaged across the palatal shelf shown for each specimen from which illustrated in situ is taken for gene of interest (coloured trace) and Shh (black trace) for the epithelium and mesenchyme. Scalebars = 200 μm.

Although each of the pathway ligands is casually referred to in the literature as a morphogen, in the RD sense any of these could in fact be a uniform, permissive component of the system rather part of the periodicity generating network. To be an RD-type patterning morphogen the activity of its pathway must also be periodic. To determine where each of these pathways is active, we analysed well-established direct transcriptional targets of their respective transduction pathways (Figs. 1B). Using pathway targets as the operational measure of pathway activity avoids the complications of multiple ligands, receptors and transduction components as discussed above. We found that each of the pathways had periodic outputs. Where available, additional target markers were tested and gave the same periodic pattern (**Fig.1 supplement 3**). As expected, Shh target expression was in the same spatial phase as Shh in both epithelium and mesenchyme. Surprisingly, FGF signalling was in-phase with the Shh stripes in the mesenchyme but out-of-phase in the epithelium. This reconciles our previous description of FGF activity as being in phase with the Shh stripes (Economou et al., 2012) with a previous report describing it as out-of-phase (Porntaveetus et al., 2010). It also means that FGF signalling must be functioning as effectively two different pathways. We refer to these pathways as mesenchymal FGF (m-FGF) and epithelial FGF (e-FGF), (although these should not be assumed to be two specific individual ligands but more likely to be combinations of ligands working through the two relatively localised receptors (Porntaveetus et al., 2010)). Wnt target expression was in-phase in the epithelium and undetectable in the mesenchyme, while BMP signalling was out of phase in both layers (Fig.1b). In summary, Shh, m-FGF and Wnt (pathways) are in-phase with the rugae, and e-FGF and BMP (pathways) are out-of-phase. The above findings show that each of the pathways is periodic and is therefore, by definition, part of a periodic pattern-generating network.

We then asked how four or potentially five components can be wired together in a regulatory network that can generate the observed spatial pattern. One potentially simplifying approach is to ask whether the topology of this system (i.e. the network – technically a directed graph – where chemical components are linked by activation or inhibition arrows) could effectively be described as a classical two-component RD system, which would consist of two “master morphogens” with the other components serving as intermediates (so-called “mediators” (Cho et al., 2011)) between them. Any pair of morphogens could potentially support a two-component system (if “in phase” by a classical Activator-Inhibitor (AI), if “out-of-phase” by a Substrate-Depletion (SD) configuration). We therefore turned to perturbation analysis as a way of constraining possible network topologies. We modelled the effects of perturbing either component in both classical two-component AI and SD systems **(Fig.2a)**. We then compared the results with the effects on established spatial patterns of experimentally inhibiting Wnt, BMP or Hh itself, considering first all four possible two-component systems containing the Hh pathway (our readout) and excluding FGF (where two nodes are inhibited simultaneously requiring a different analysis – see below). We found that for both Wnt-Hh and BMP-Hh systems, one of the two possible two-component models was consistent with the data (Fig. 2b). The two topologies cannot be combined into a three-component system in which one of the components can be collapsed away (as it were, embedded) as a passive mediator of a two-component master pair (Fig. 2 and Fig 2 supplement 2). This showed that our system (probably like most real systems) can *not* be accurately described as falling into either one of the two classical AI or SD classes.

**Fig 2.**
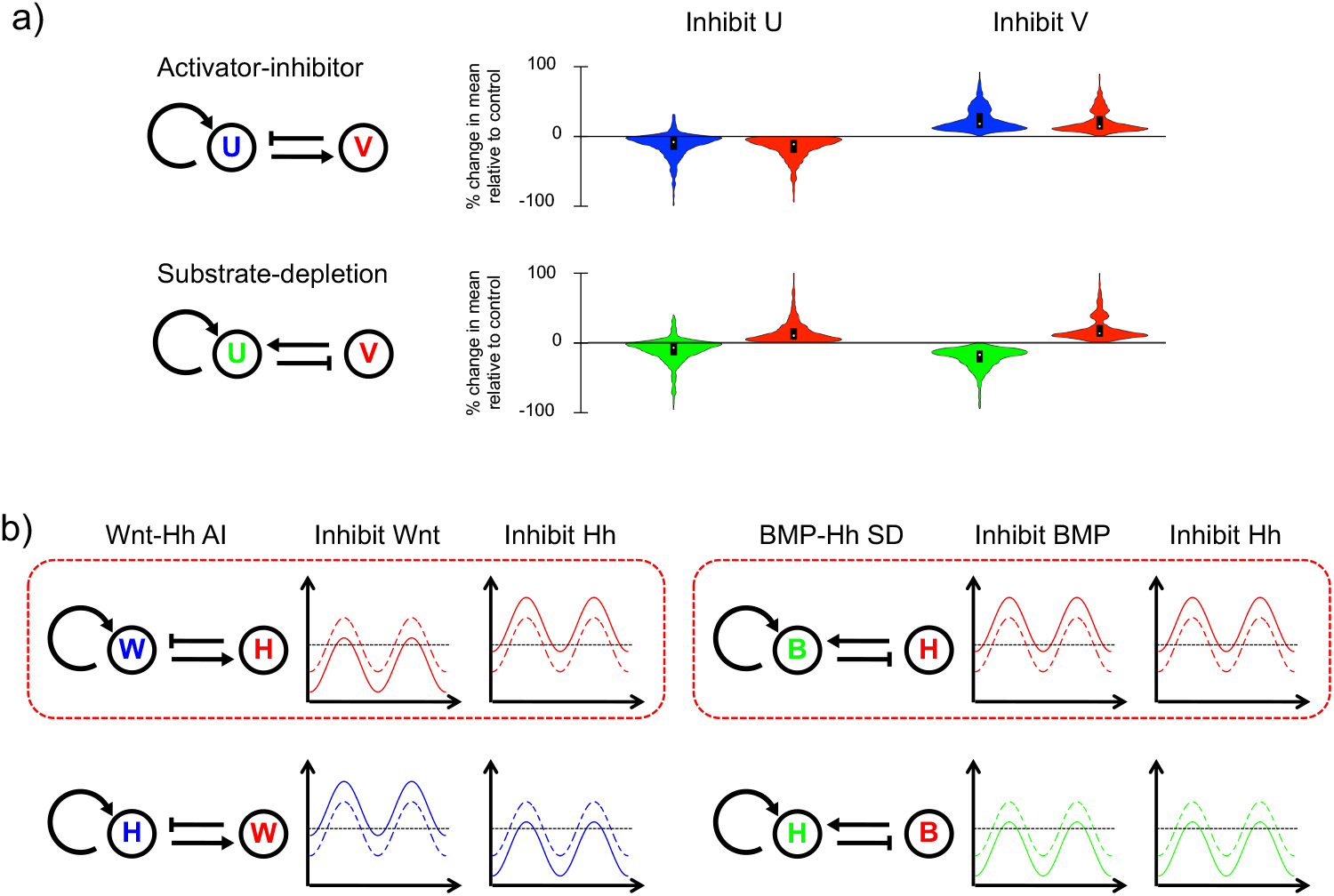
Numerical simulation of reaction-diffusion patterning inhibition for two-component systems. a) Violin plots showing percentage change in the mean level of components U and V in illustrated activator-inhibitor (AI) and substrate-depletion (SD) RD networks, on inhibition of the response to component U and V in RD simulations (plots for inhibition of response shown in fig 2 supplement 1). b) Networks showing the two possible configurations of Wnt-Hh AI systems and BMP-Hh SD systems, with components coloured according to equivalent component in a). Next to each is a schematic of the response of Hh on inhibition of each component in the network, based on the predominant response in the simulations. Solid lines indicated levels after inhibition, with dashed lines representing uninhibited states. The two topologies whose responses to inhibition replicate the experimental observations (see fig 1 a) are highlighted with a red box.

The above results showed that modelling this system requires consideration of higher order (i.e. greater than two-component) networks. We therefore next asked whether any three-component networks could be found consistent with the spatial pattern and perturbation data. There are 3^9^ = 19,683 possible 3-component network topologies, since each component can interact with the two others and itself (nine types of interaction) and each interaction can be positive, negative or zero (**Fig. 3 supplement 1**). Considering that all components have some degradation rate, and therefore have some level of negative self-interaction, we only considered the distinction between topologies with positive self-interactions and those without them. This reduces the total number of possible topologies to 2^3^ × 3^6^ = 5,832. (These different topologies incidentally place different constraints on diffusion, as discussed in (Marcon et al., 2016)).

To go further, we resorted to numerical methods because, unlike for the two-component system, there is not a well-established relationship between network topology and spatial patterning for 3-component networks (Scholes, Schnoerr, Isalan, & Stumpf, 2019). Three-component RD systems were considered by White and Gilligan in the context of hosts, parasites and hyper-parasites (White & Gilligan, 1998), where they described criteria that determine which such systems generate DDI, as well as criteria to distinguish temporally stable from oscillating systems. More recently, related analyses, including graph-based approaches, have been applied to developmental periodic patterning (Marcon et al., 2016) and general RD systems (Diego, Marcon, Müller, & Sharpe, 2018; Scholes et al., 2019). We systematically screened parameter ranges around known values published for biological RD systems (Nakamasu, Takahashi, Kanbe, & Kondo, 2009) and applied the White and Gilligan (White & Gilligan, 1998) criteria for DDI in a 3-component model. Out of 242,121,642 parameter sets (9 interactions × 7 values for each × 6 different choices for which morphogens have high or low diffusion values) searched we found a subset of 653,574 parameter sets that gave DDI. The signs of the parameters define specific topologies. As the rugae form as stable stripes of gene expression and tissue thickening (Economou, Brock, Cobourne, & Green, 2013; Economou et al., 2012), our first observational constraint was that the stripes we see are non-oscillating. We recovered 1,492 topologies giving non-oscillating DDI (see Methods for details). A second constraint was that inhibition of any of the components would perturb the pattern. This implied that every node (morphogen) in the network outputs as well as inputs making the network “strongly connected” (Marcon et al., 2016). We recovered 1,396 strongly connected network topologies. Among these, the four possible phase relationships (all-in-phase or each one of the three out-of-phase with the other two) could each be generated by 498 different topologies (with some topologies capable of generating more than one possible phase relationship depending on parameter values).

To investigate the 498 topologies in a given phase group, we generated “stalactite” plots according to Cotterell and Sharpe (Cotterell & Sharpe, 2010). In this representation, each topology is shown as a point linked to all other topologies that differ from it by the gain/loss of one edge, with each row corresponding to the total number of edges. Topologies at the base of each stalactite, of which there were 45 for each phase group, represent those that are distinct (non-overlapping) and possess only non-redundant interactions (Fig.3A).

**Fig 3.**
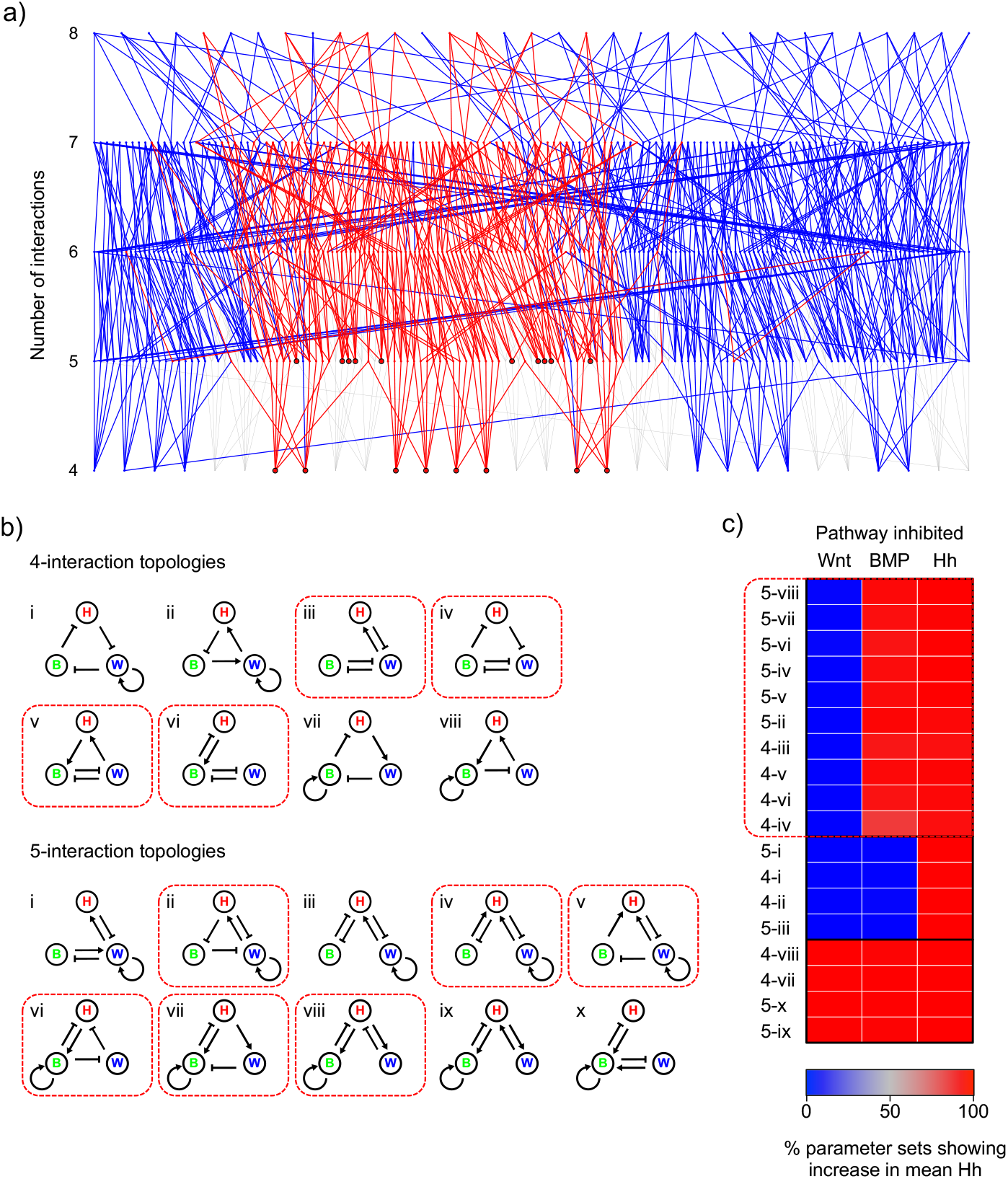
“Stalactite plot” and numerical simulation identifying subsets of 3-component RD systems and their behaviours under inhibition. a) Topology atlas for the phase group of 3-component networks identified in the parameter search, consistent with the spatial pattern of Wnt, BMP and Hh. Each node represents a topology. Those that are not “strongly-connected” are greyed out; those with fast-diffusing Hh and slow diffusing Wnt and BMP are in red. Of the 45 topologies at the bottom of stalactites, the 18 that are consistent with the diffusion constraints are outlined in black. b) The 18 Wnt-BMP-Hh 4- and 5-interaction networks. Within each group networks are numbered according to their position from left to right in a). c) Heat map showing the percentage of parameter sets where the level of Hh increases in response to the inhibition of each component in the network in RD simulations. Topologies are arranged by hierarchical clustering. The 10 topologies whose response is consistent with the experimental data are highlighted with a red box, as are the network diagrams in b).

We could now begin to consider real morphogens as nodes in the identified topologies, to ask which topologies could correspond firstly to the observed spatial phase relationships and secondly to likely diffusivities. We considered three-node networks consisting of Shh, Wnt and BMP. We knew the phase relationship of these from direct observation of the targets shown in Fig.1: Shh and Wnt activities are in-phase with the rugae and BMP activity is out-of-phase. This constraint identified a specific phase group as of 45 topologies as relevant to our system. As for diffusivities, we observed that the peak-widths of Shh target gene expression were consistently somewhat larger than those of the other components and of its ligand (Fig.1b) making it likely that Shh was the fast-diffusing component compared to the other two. (We show below that this provisional assumption is ultimately not required for selection of networks consistent with experiment.) These phase and diffusivity constraints reduced the number of working topologies down to just 18 (Fig.3a,b).

With these 18 topologies, we now compared their predicted behaviour under perturbation to the experimental results using numerical simulations similar to those described above (see Methods for details). The numerical modelling revealed three groups of characteristic responses to inhibitions (Fig.3c and Fig.3 supplement 2). Ten out of the eighteen networks showed Hh pathway responses consistent with the experimentally observed changes in Shh expression (boxed in Figure 3c).

From several thousand conceivable networks, the above analytical, numerical and experimental methods identified just ten three-component, minimally connected topologies that captured the experimentally observed periodicity, phase and perturbation responses of three of the five components identified as active in this system. We will refer to these as “valid” networks. How might this help us discover what networks would be valid that include all five components, and potentially “n” additional components?

It has been suggested that general rules for biological circuit behaviour can be inferred from analysis of the signs of the embedded feedback loops (Tyson & Novak, 2010). We therefore examined whether this approach might be applied to our RD networks. We observed that our valid three-component networks contained the same feedback loops as the two perturbation-consistent 2-component networks (Fig. 2B). In the 4-interaction networks, Wnt and BMP are in positive feedback loops *by mutual inhibition* while Hh is always only in a negative feedback loop (Wnt and BMP can also be found in the negative feedback loop), while in the 5-interaction networks, Wnt and BMP are still in a positive feedback loop through (direct or indirect) mutual inhibition, while the remaining feedback loops of either 2-component system are embedded in the network (Fig. 3B and Fig. 3 – supplement 3 indicating the loops). In other words, the behaviour of the network, including under perturbation, is embedded in the product of the signs of the “arrows”, i.e. the signs of the reaction term coefficients. We investigated numerically whether this is the case and found (Supplementary Note 3A) that for all but very small perturbations (in which diffusion can dominate) or very large perturbations (in which RD breaks down altogether), that it is indeed possible to predict the behaviour of an RD-competent network from its reaction terms alone. Analytical considerations (Supplementary Note 3B) reinforce this conclusion.

The analysis revealed that the relationship between reaction terms and the behaviour of components in response to perturbation has a consistent mathematical form (asymptotic curves). This enables the sign of the perturbation response (as opposed to its precise magnitude) to be predicted as a relatively simple set of conditions fulfilling or not fulfilling certain inequalities (Supplementary Note 2). We were able to express the terms in the inequalities as functions of the combination of signs of the reaction terms (i.e. feedback loops) in n-component systems, thus predicting the effects on perturbation of any given RD system component depending on its participation in positive and/or negative feedback loops (Supplementary Note 3). We were able to use this approach to analyse how the response of components to inhibition in the system changes upon step-wise addition of new nodes (components) (Supplementary Note 4). In brief, an RD system requires that for an n-node system, there needs to be a positive feedback loop through a maximum of n-m nodes and a negative feedback loop through the remaining m nodes (provided n > m ≥ 1) and at least one node must be in both loops. Examples of loops in 2-node (classical RD) and 3-node systems are shown in Fig. 4A. With these conditions in mind, one can predict outcomes of perturbations of each type of node included in these loops (Fig. 4B). However, the above conditions allow for networks where the loops go through only a subset of nodes. The predictions of outcomes of perturbations in Fig.4B therefore need not apply. Further analysis (decribed in Suplementary Note 4) generated predictions of what happens to components represented by nodes outside the RD-defining “core” loops (Fig. 4C,D). We found that effects of inhibition of these “non-core” components to be mostly similar, but not identical, to those of inhibition of core components. (This means that sometimes alternative “cores” in a given network will produce different perturbation responses – see Discussion below)

**Fig 4.**
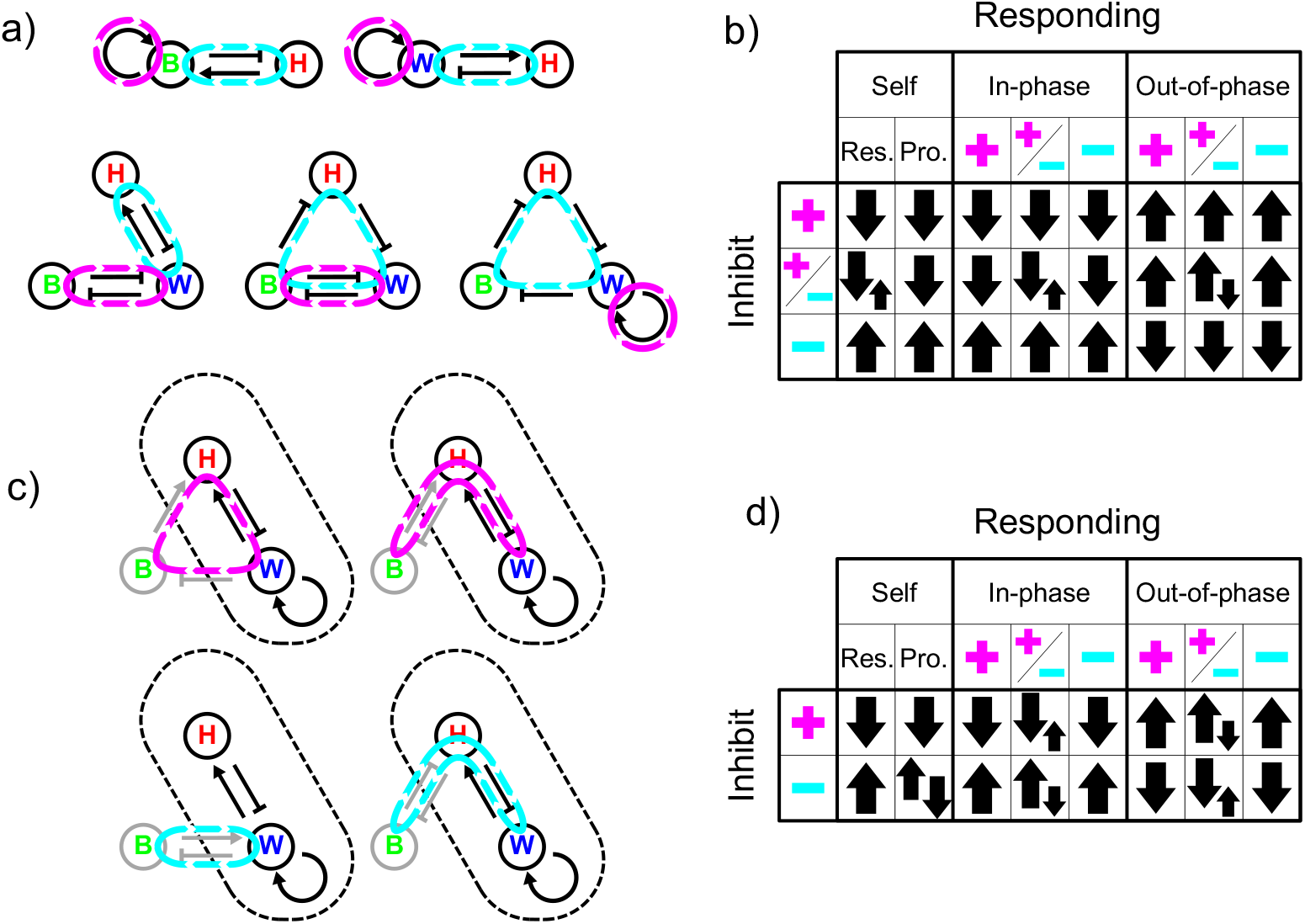
Identification of feedback loops and resulting behaviours under inhibition for 3-component RD systems. a) Illustrative 2- and 3-component networks showing minimal feedback requirements for RD of one or more components forming a single positive feedback loop (highlighted in magenta), with a negative feedback loop being formed through at least one addition component (highlighted in cyan). b) Summary constraint table showing response of components in such a minimal network to the inhibition of a component found in either the positive feedback loop alone (magenta ‘plus’ sign), negative feedback loop alone (cyan ‘minus’ sign) or both (‘plus’ and ‘minus’), depending on which loop they are in and the phase relative to the inhibited component (‘In-phase’ or ‘Out-of-phase’). For the response of a component to its own inhibition (‘Self’), the inhibition of response (‘Res.’) and production (‘Pro.’) are shown. Up arrows indicate an increase in a component’s levels, down arrows a decrease. Two arrows of equal size show when the system is unconstrained. Where opposing large and small arrows are shown, the system behaves according to the large arrows, apart from under certain topologies (see supplementary note 4) where the reverse is seen for certain components. While components are not constrained, different components showing the unconstrained response are coupled to one another. c) Illustrative examples showing how additional components can be integrated into a minimal RD system outside of the ‘core’ RD network by forming external loops. External components and interactions are shown in grey either forming a positive or negative feedback loops with the core positive feedback loop (highlighted in magenta and cyan respectively). Core RD network outlined with dashes. d) Summary constraint table showing response of components in such a network to the inhibition of a component either providing additional in the positive feedback (magenta ‘plus’ sign) or negative feedback (cyan ‘minus’ sign) to the core positive feedback loop, depending on which loop in the core network they are in and the phase relative to the inhibited component. Symbols as in b). For subset of topologies where a component is in both loops see supplementary note 4.

We applied these rules for perturbation responses and effects of node addition to the 45 Wnt-BMP-Shh networks discussed above that fulfilled the requirements of diffusion-driven instability and the observed phase relationships. Systematically identifying all possible loop combinations (Fig. 4 supplement 1) enabled prediction of the responses of each component to any inhibition using the constraint tables in Fig. 4. This showed that the observed perturbation responses corresponded to the same ten topologies as in Fig. 3 but in this case without the prior assumption that Shh was the fast-diffusing component.

The effect of inhibition of FGF can now be reconsidered. Since the effect on Shh was neither net increase nor net decrease, we considered topologies in which the eFGF and mFGF acted in opposition. Simulation with a parameterisation of the very simple network of this kind depicted in Fig. 5A showed that a blended increase and decrease leading to a flatter waveform was indeed obtained (Fig. 5B). Thus, a RD system with two FGFs, each having opposite effects on Shh, can account for our experimental observations.

**Fig 5.**
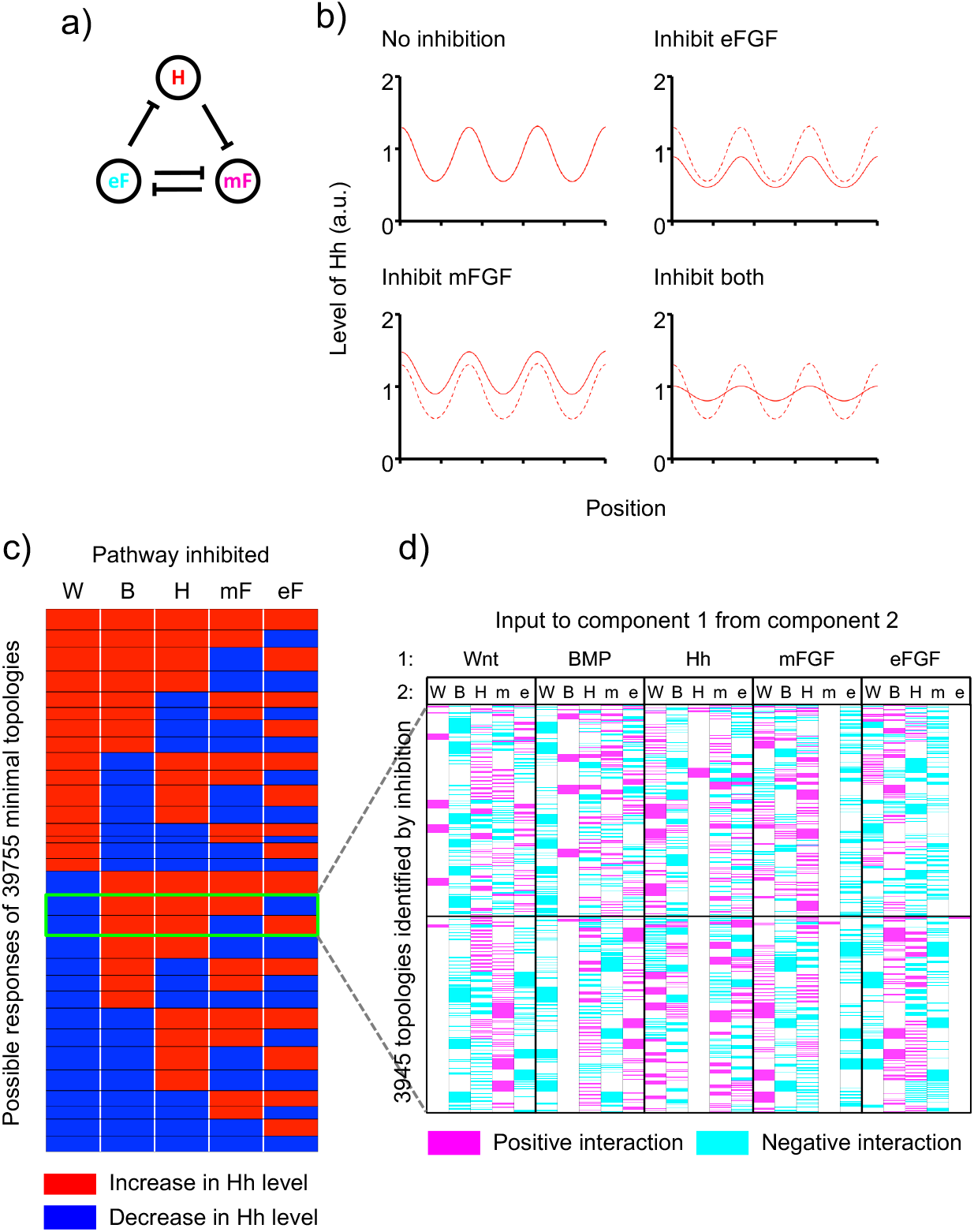
Integration of two out-of-phase FGF morphogens explains FGF-inhibition effects and allows prediction of behaviours under inhibition of 5-component RD systems. a) Example of a network where the inhibition of epithelial FGF (eF) and mesenchymal FGF (mF) would be predicted to have opposing effects on the levels of Hh (H) according to analysis of reaction terms. b) Simulation of single and combined inhibition of the two FGF components (dashed lines indicate uninhibited state). Simulations carried out as detailed in methods with u_1_ is mF, u_2_ is eF and u_3_ is H, with *a_12_* = −0.019, *a_13_* = −0.034, *a_21_* = −0.019, *a_32_* = −0.022, *b_1_* = 0.064, *b_2_* = 0.037, *b_3_* = 0.068, *c_1_* = 0.004, *c_2_* = 0.011, *c_3_* = 0.039, *fmax_1_* = 0.008, *fmax_2_* = 0.022, *fmax_3_* = 0.072, *D_1_* = 1.03, *D_2_* = 1.71 and *D_3_* = 7.63. Where not specified a_ii_ = 0. c) Map of all possible responses of Hh to inhibition of each of the five components for all 39755 predicted minimal topologies (red = increase, blue = decrease) arranged into 31 groups of responses, outlined in black. Two groups (3945 topologies) matching the observed responses (Wnt down, BMP up, Hh up, mFGF and eFGF opposing responses) highlighted in green box. d) Map showing the interactions making up the 3945 topologies identified by perturbation analysis (positive interactions in magenta, negative in cyan, no interaction in white). Horizontal black line separates topologies giving different patterns of responses of Hh in response to mFGF and eFGF inhibition (green-boxed groups in c).

We considered all possible 5-component topologies with loops required for DDI with the sign of links constrained by their phase relationship (see Supplementary Note 4). Specifically, we first identified all possible sets of core topologies (i.e. all sub-networks of two to five components that have the minimal number of interactions sufficient for RD). For each core, all possible combinations for wiring-in the remaining components were recovered (code available in Supplementary materials). This yielded 39,755 unique minimal topologies. Prediction of their responses to inhibition revealed that, of these, 3,945 were consistent with our experimental perturbation results. Fig. 5D depicts these topologies in terms of the interactions (arrows) between the five components. While some interactions are different in different topologies, some are the same or absent for all topologies (e.g. the input of mFGF to eFGF indicated in the second-to-right most column are inhibitory (cyan) or absent (white) in all topologies).

What further experimental data could constrain potential network topologies? We turned to temporal behaviour of the rugal system. We conducted a large number of in situ hybridisations on adjacent sections from a long time-series of specimens, spatially registering multiple specimens to the stripes of Shh expression. Because the pattern arises through monotonic linear growth (Economou et al., 2012), we could temporally order and space our specimens. This allowed us to determine the kinetics of the onset of stripe formation for each of the pathway markers (Fig. 6).

**Fig 6.**
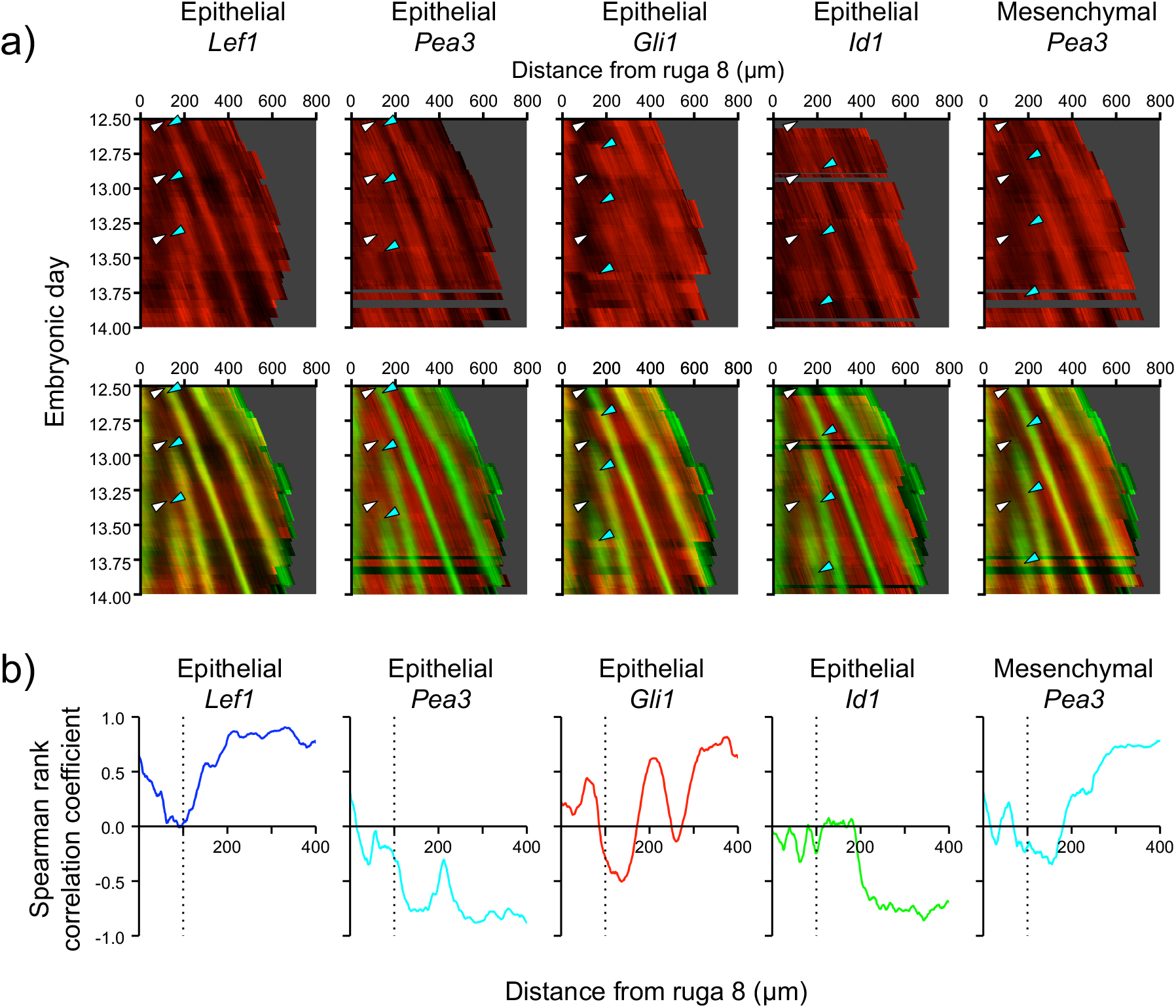
Periodic gene expression kymographs reveal an early Wnt-epithelial-FGF-Shh initiating “core” system with mesenchymal FGF and BMP integrated later. a) Kymographs showing the pattern of expression of indicated targets genes through time (red) and their expression relative to *Shh* (green) for rugae 3, 4 and 5. White arrowheads indicate the onset of *Shh* expression for each ruga, and cyan arrowheads indicate the change in the expression pattern of each target gene associated with each ruga. Mesenchymal *Gli1* and *Id1* expression resemble the epithelial patterns (data not shown). b) Plot of the Spearman rank correlation coefficient for the intensity of *Shh* staining and the marked target gene across all time points for each position relative to ruga 8, indicating when relative to the onset of *Shh* expression the spatial pattern for each target gene emerges. Dotted line at 100 μm indicates the onset of *Shh* expression.

Expression of the Wnt target *Lef1* and loss of FGF target *Pea3* in the epithelium are simultaneous with the onset of *Shh* transcription (Fig.6). On the other hand, downregulation of BMP target *Id1* and the increase of FGF target *Pea3* in the mesenchyme lag by approximately six hours (Fig. 6). This lag is most consistent with a 3-component “core” network consisting of Wnt, e-FGF, and Hh signalling initiating periodic stripes, with BMP and m-FGF subsequently incorporated into the network.

Plotting correlation of other markers in relation to *Shh* expression provides further evidence that *Shh* is indeed the inhibitory component of the network. The upregulation of *Lef1* and the downregulation of *Pea3* are essentially concomitant with the onset of expression of *Shh* but they occur where *Shh* signalling (i.e. *Gli1* expression) is low (i.e. inhibition is low). The response to *Shh* is evident at a slight delay, which is observed as the trajectory of inhibition in all RD simulations.

What does the dynamic data mean for our networks? As e-FGF and Wnt responses are the first movers and move simultaneously, they must form a core positive feedback loop together with Hh providing the negative feedback loop necessary for RD. As BMP and m-FGF responses trail Hh, they cannot be part of the core positive feedback loop which destabilises the system. Applying these constraints to the 3,945 topologies that were consistent with the experimentally observed phase relations and responses to perturbation gave 154 topologies consistent with the observed dynamics (0.004 % of the total 39,755 DDI minimal topologies) (Fig.7). All of these are subsets of the same set of regulatory interactions (Fig.7c), comprising one of four possible Wnt-eFGF-Hh cores, with BMP and m-FGF providing additional interactions.

**Fig 7.**
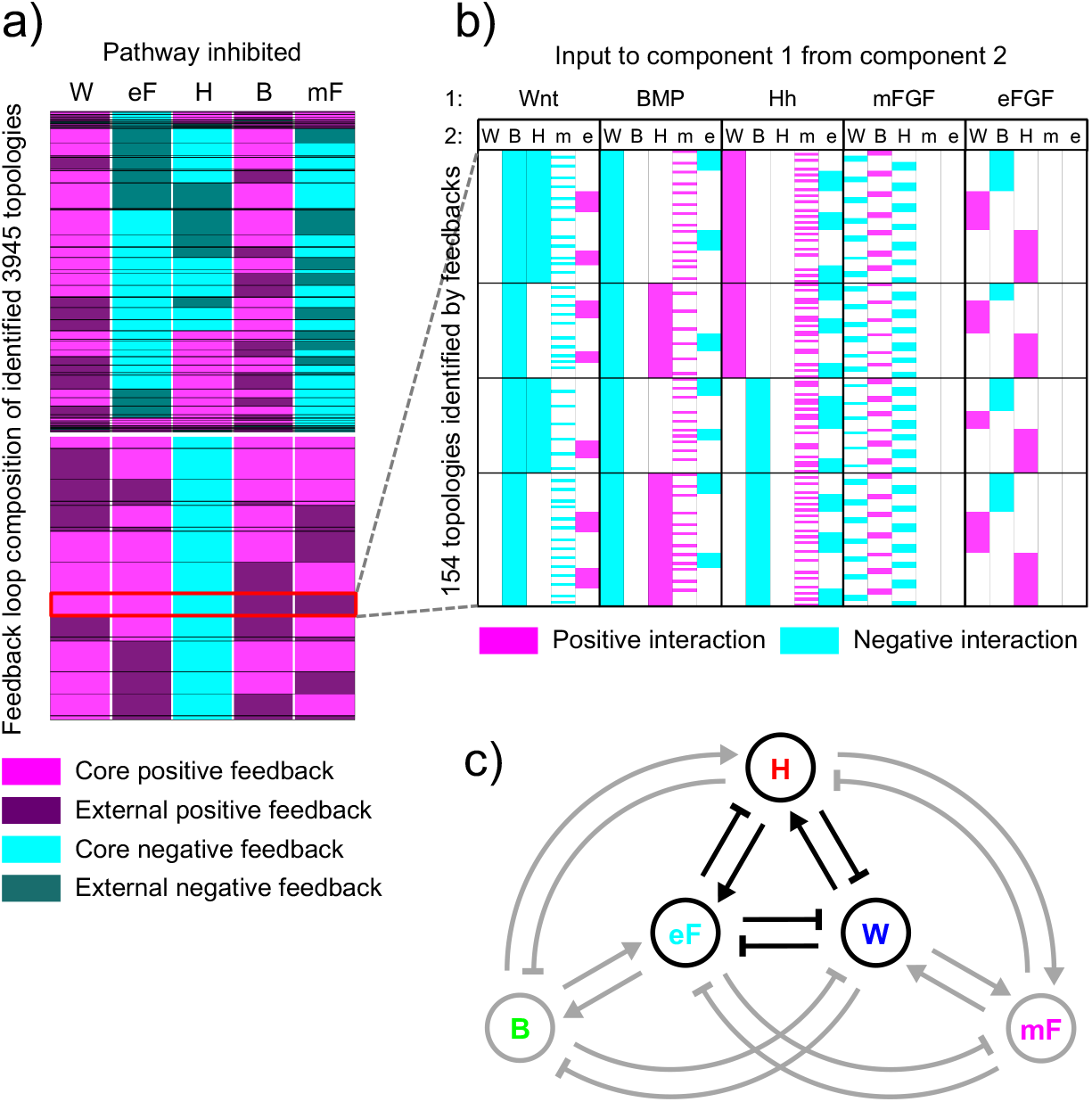
Possible feedback loops and network topologies constrained by experiment. a) Map showing feedback loop structure of 3,945 topologies by perturbation analysis (see fig 5c and d). Components involved in positive feedback in magenta, and negative feedback in cyan, with components external to the core in dark. Horizontalwhite line separates topologies giving different patterns of responses of Hh in response to mFGF and eFGF inhibition (upper group of topologies correspond to upper group of highlighted topologies in figs 5c and d). Groups of topologies showing different patterns of feedback loops are outlined in black. Set of 154 topologies showing constraints on topology as determined by kinetic analysis (Wnt and eFGF as only core positive feedback components and Hh as core negative feedback component) highlighted in red. b) Map showing the interactions making up the 154 topologies identified by feedback loop analysis (positive interactions in magenta, negative in cyan, no interaction in white). Horizontal black lines separate four groups of topologies with different interactions between the three core components Wnt, eFGF and Hh. c) The 154 topologies identified in a) and shown in b) all show the same signs of interactions where present. Summary network diagram showing the signs of the interactions found among these topologies.

## Discussion

We have shown that five classical morphogen pathways periodically pattern the rugae and have identified a relatively small number of potential Reaction-Diffusion network topologies and an even smaller number of consistent regulatory interactions between nodes within such networks. The modelling, combined with the experimentation thus far, suggests – although of course does not prove – potentially direct molecular interactions between specific pathway-induced transcription factors and target enhancer sites in the genes encoding the other morphogens.

This study highlights more generally the ways in which experimental data can be used to challenge RD models in a mammalian tissue context. In particular, we have shown that the effects of relatively small acute perturbations to spatial patterns that have already formed are highly constraining on plausible network topologies. This suggests that similar inhibitor studies can be a useful complement to “knockout” genetics in understanding these dynamical systems. We also gained constraining information from the dynamic (temporal) evolution of the system during embryonic development. This was facilitated by the sequential appearance of the stripes, which is a peculiarity of this rugal palate system. In many periodic patterns, such as, for example, the appearance of cartilage ring-patterning stripes in the developing trachea (Sala et al., 2011), stripes appear simultaneously and so identifying leading and lagging genes would require more quantitative measurements.

It is striking that component number (even at arbitrarily chosen level) reduces constraint very rapidly. Notably, at 3-component level there is a small part of parameter space for which networks give behaviours that do not conform to the minimal topologies from which they are derived. How this grows and whether it could be important at higher orders is not clear. However, the “black boxing” of whole pathways into single nodes provides a way of rationally reducing the system complexity in a way that makes biological sense. It remains an open question as to how to integrate this way of thinking with ‘omic datasets. There is a significant body of work on analysis and simplification of complex regulatory networks and some thought has been given to RD processes within these (Mincheva & Roussel, 2012). However, practical application of these methods with experimental input is still in its infancy.

Apart from the computational aspects of our work, it raises biological questions. Why, for example, does the patterning of the palate use five pathways when, in principle, two would do? One possibility is that this provides a particular type of robustness to perturbations: yes, there are many targets whose mutation can affect the pattern, but there is also significant redundancy such that the modifications to the pattern are mostly relatively subtle. Another possibility is that each pathway provides an additional tuning of the pattern or setting up of signalling for downstream events such as differentiation. A completely opposite explanation is that all of the regulatory interactions exist in cells and are used elsewhere for multiple other purposes (for RD, or not and in pairs or not) and that the apparent “overkill” in terms of numbers of pathways involved is merely because there is no evolutionary pressure to eliminate or suppress their role in the palate. More detailed analysis of the robustness properties of these networks and of the conservation of the regulatory interactions is needed to address these questions.

## Supporting information

Supplementary Figs (all)

## Acknowledgements

Atsushi Ohazama, Martyn Cobourne, Abigail Tucker, Albert Basson and David Rice for gifts of in situ probe constructs. This work was funded by BBSRC grant BB/J009105/1 to J.B.A.G. We thank Attila Czikas-Nagy and Ton Coolen for critical reading of the manuscript.

