## Supplementary Figs (all) for "Perturbation analysis of a multi-morphogen Turing Reaction-Diffusion stripe patterning system reveals key regulatory interactions"

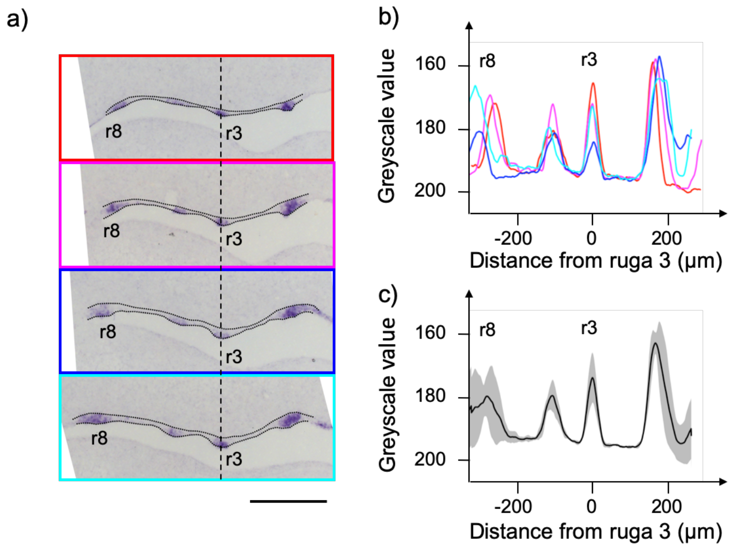


**Figure 1 supplement 1 Quantification of striped rugal expression**

a) In situ hybridisation showing *Shh* expression in sagittal sections taken at 35 µm increments across a E13.5 palatal shelf, ordered from medial most (red outline) to lateral most (cyan outline). The positions of ruga 3 and ruga 8 are indicated. Dotted lines illustrate the extent of the palatal epithelium used for quantifications. Anterior to right. Scalebar = 200 µm.

b) Intensity profiles for each section aligned by the position of ruga 3. Peaks in *Shh* intensity at ruga 3 and ruga 8 are indicated. Profiles coloured as in a).

c) Mean intensity profile from the four sections in a) and b). Shaded area represents 1 sd.

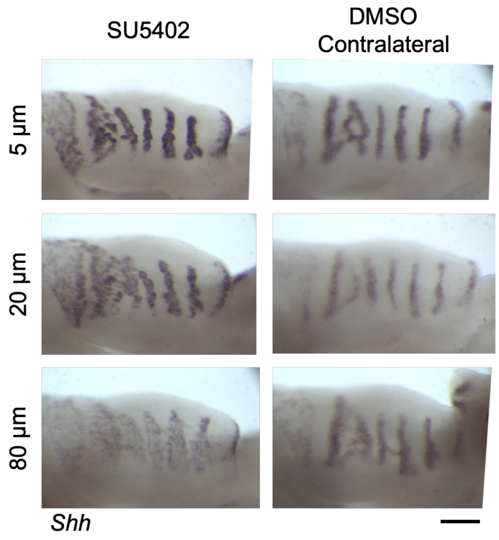


**Figure 1 supplement 2 Titration of FGF inhibitor SU5402**

Shh expression in palatal shelves explanted at E13.5 and cultured with the FGF receptor inhibitor SU5402 for 24 hrs at the indicated doses, alongside the contralateral shelves which were cultured in DMSO. Scalebar = 200 µm.

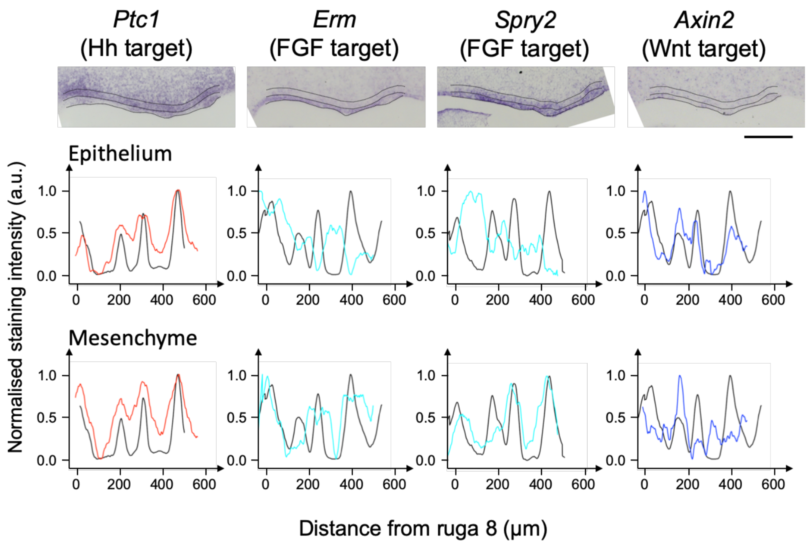


**Figure 1 supplement 3 Additional pathway marker quantification**

In situ hybridisation of sagittal section through E13.5 palatal shelf for additional markers of pathway activity. An additional marker of BMP signalling (*Id2*) was tested but did not show specific staining in the palate (data not shown). Dotted lines illustrate the extent of the palatal epithelium and the underlying mesenchyme used for quantifications. Anterior to right. The intensity profile averaged across the palatal shelf shown for each specimen from which illustrated in situ is taken for gene of interest (coloured trace) and Shh (black trace) for the epithelium and mesenchyme. Scalebars = 200 µm.

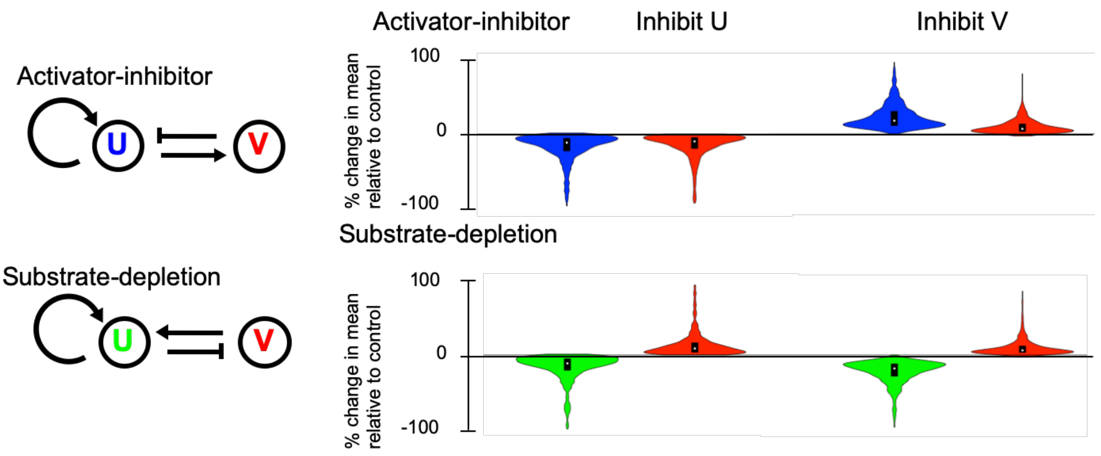


**Figure 2 supplement 1 Inhibition of production produces the same outcomes as inhibition of response**
Violin plots showing percentage change in the mean level of components U and V in illustrated activator-inhibitor (AI) and substrate-depletion (SD) RD networks, on inhibition of the production of component U and V in RD simulations.

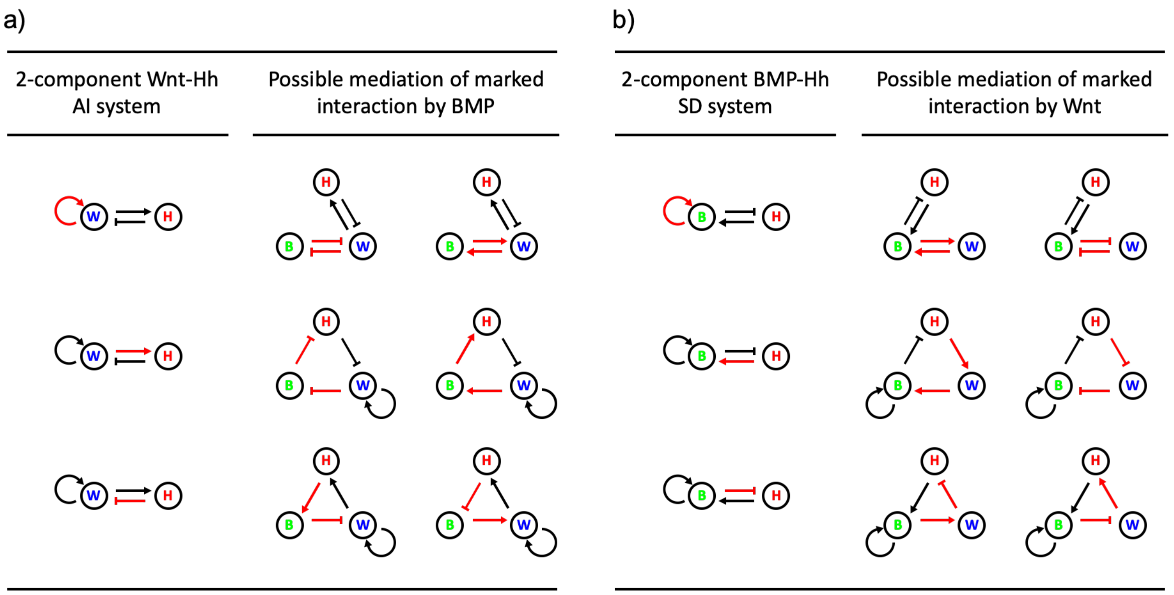


**Figure 2 supplement 2 Adding a component as a mediator of an interaction**
a) Possible ways that an interaction in a two-component Wnt-Hh AI network (marked in red) can be 'mediated' by BMP in the adjacent 3-component Wnt-BMP-Hh RD networks. Interaction in the 3-component RD networks maintain the net sign of the marker interactions in the 2-component networks; the phase of BMP relative to Wnt and Hh is not considered.
b) Same as in a) for Wnt mediating interaction in a 2-component BMP-Hh SD network.
Wnt in blue, BMP in green and Hh in red.

a)

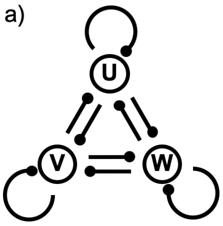


b)

| | |
|---|---|
| All possible topologies: | **19683** |
| Degradation by default: | **5832** |
| Stable DDI topologies: | **1492** |
| Strongly connected: | **1396** |
| For each phase relationship: | **498** |
| Minimal topologies (per phase): | **45** |

**Figure 3 supplement 1: Three-component topologies and exclusion by successive constraints**

a) 3-component network topology showing the nine possible interactions captured in the reaction matrix.

b) Enumeration of topologies recovered from parameter search under different constraints as detailed in text.

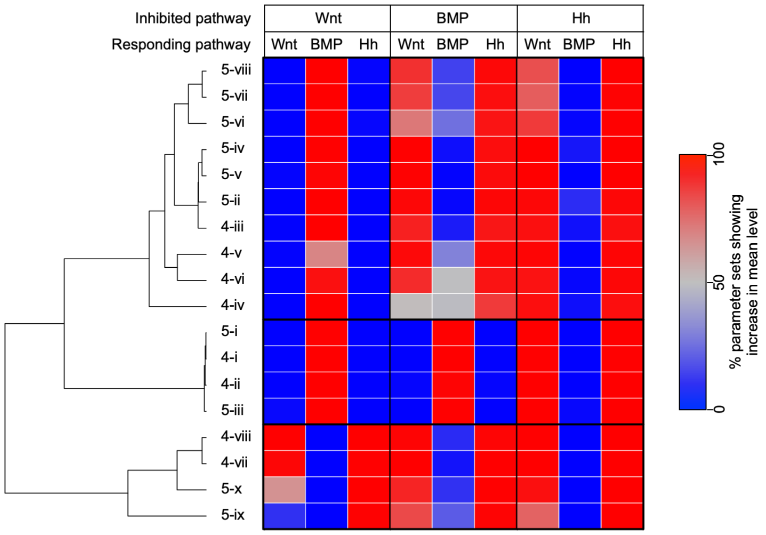


**Figure 3 supplement 2: Responses of 3-component networks to perturbation**

Full heat map showing the percentage of parameter sets where the level of each component increases in response to the inhibition of each component in the network in RD simulations. Topology names are as in figure 3. Hierarchical clustering shows that the behaviours of the topologies fall into the main categories for which all three components show similar behaviour.

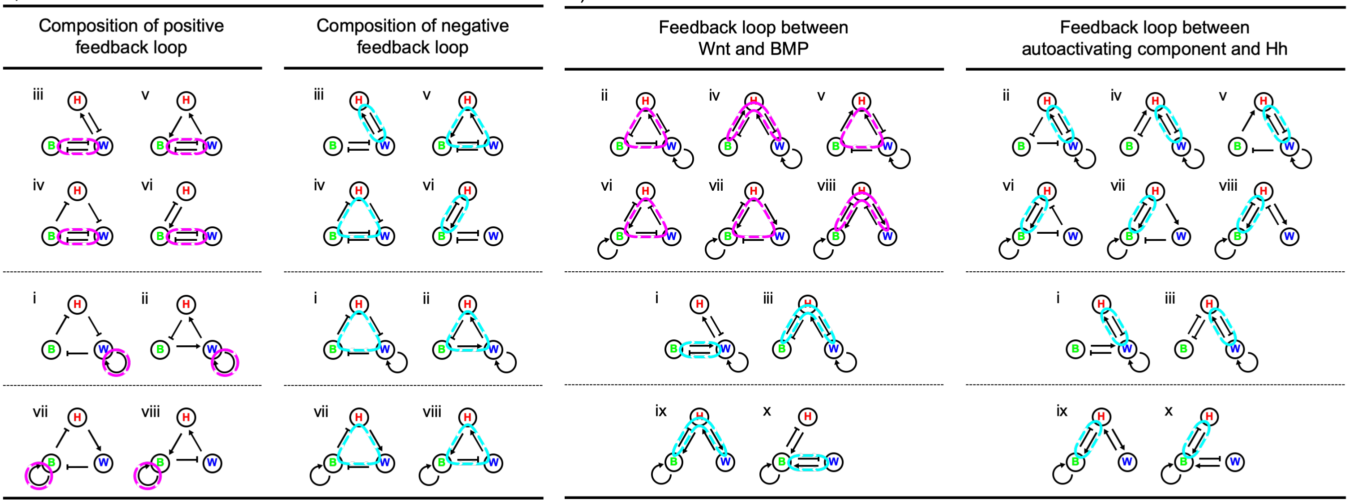


**Figure 3 supplement 3: Feedback loops in 3-component networks**

a) Schematics showing the eight minimal 4-interaction RD networks recovered from the parameter search, with either the single positive feedback loop marked in magenta (left) or the single negative feedback loop marked in cyan (right). Topologies are group as according the the hierarchical clustering in figure S7.

b) Schematics showing the ten minimal 5-interaction topologies networks recovered from the parameter search. On left, the net feedback loop between Wnt and BMP is marked, with magenta for a positive feedback, or cyan for a negative feedback. On the right, the presence of a negative feedback between Hh and the autoactivating component (Wnt or BMP) is marked in cyan. Topologies are group as according the the hierarchical clustering in figure S7.

Wnt in blue, BMP in green and Hh in red.

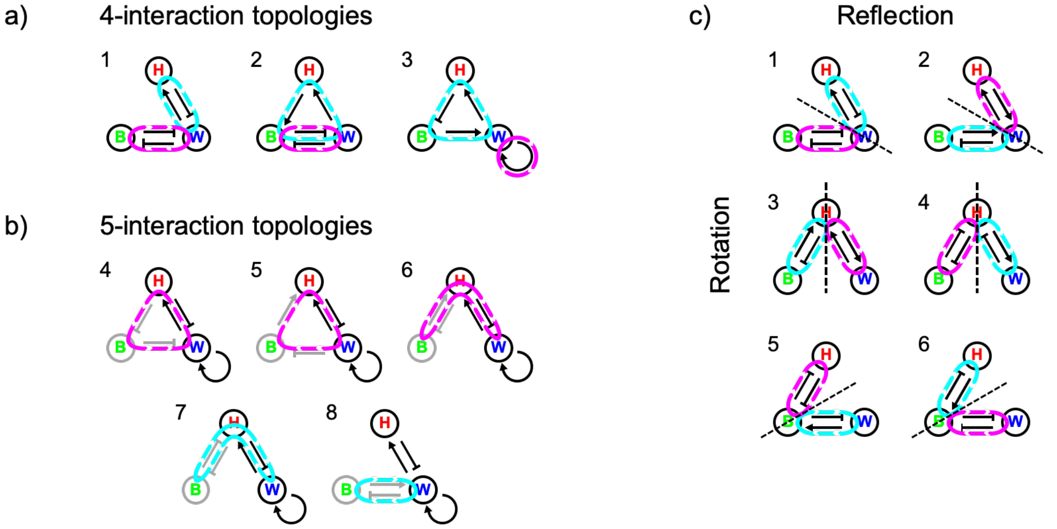


d) Response of Hh

| | Topology | 3 | 3 | 7 | 8 | 7 | 2 | 6 | 5 | 1 | 4 | 2 | 6 | 5 | 1 | 4 | 7 | 3 | 8 | 3 | 7 |
|---|---|---|---|---|---|---|---|---|---|---|---|---|---|---|---|---|---|---|---|---|---|
| | Orientation | 6 | 5 | 5 | 5<br>6 | 6 | 5 | 5 | 5 | 5 | 5 | 4 | 4 | 4 | 4 | 4 | 2 | 2 | 1<br>2 | 1 | 1 |
| Inhibited pathway | Wnt | ▲ | ▲ | ▲ | ▲ | ▲ | ▲ | ▲ | ▲ | ▲ | ▲ | ▲ | ▲ | ▲ | ▲ | ▲ | ▼ | ▼ | ▼ | ▼ | ▼ |
| | BMP | ▲ | ▲ | ▲ | ▲ | ▲ | ▬ | ▲ | ▲ | ▲ | ▲ | ▲ | ▲ | ▲ | ▲ | ▲ | ▼ | ▼ | ▼ | ▼ | ▼ |
| | Hh | ▲ | ▲ | ▲ | ▲ | ▲ | ▬ | ▼ | ▼ | ▼ | ▼ | ▬ | ▬ | ▬ | ▬ | ▬ | ▲ | ▲ | ▲ | ▲ | ▲ |

| | Topology | 2 | 2 | 6 | 5 | 1 | 4 | 6 | 5 | 1 | 4 |
|---|---|---|---|---|---|---|---|---|---|---|---|
| | Orientation | 6 | 1 | 1 | 1 | 1 | 1 | 6 | 6 | 6 | 6 |
| Inhibited pathway | Wnt | ▼ | ▼ | ▼ | ▼ | ▼ | ▼ | ▼ | ▼ | ▼ | ▼ |
| | BMP | ▲ | ▲ | ▲ | ▲ | ▲ | ▲ | ▲ | ▲ | ▲ | ▲ |
| | Hh | ▲ | ▲ | ▲ | ▲ | ▲ | ▲ | ▲ | ▲ | ▲ | ▲ |

| | Topology | 7 | 3 | 3 | 7 | 8 | 2 | 2 | 6 | 5 | 1 | 4 | 6 | 5 | 1 | 4 |
|---|---|---|---|---|---|---|---|---|---|---|---|---|---|---|---|---|
| | Orientation | 4 | 3 | 4 | 3 | 3<br>4 | 3 | 2 | 2 | 2 | 2 | 2 | 3 | 3 | 3 | 3 |
| Inhibited pathway | Wnt | ▲ | ▲ | ▲ | ▲ | ▲ | ▼ | ▬ | ▼ | ▼ | ▼ | ▼ | ▼ | ▼ | ▼ | ▼ |
| | BMP | ▼ | ▼ | ▼ | ▼ | ▼ | ▼ | ▼ | ▼ | ▼ | ▼ | ▼ | ▼ | ▼ | ▼ | ▼ |
| | Hh | ▬ | ▬ | ▬ | ▬ | ▬ | ▬ | ▬ | ▼ | ▼ | ▼ | ▼ | ▬ | ▬ | ▬ | ▬ |

**Figure 4 supplement 1: Loops in 3-component networks**

a) Illustrative examples of minimal 4-interaction 3-component RD networks showing the three different ways a single positive and negative feedback loop can be combined. The positive feedback loop can pass through one (topology 3) or two (topologies 1 and 2) components, and the negative feedback can pass through two (topology 1) or three (topologies 2 and 3) components. Positive feedback loops in magenta, negative feedback loops in cyan.

b) Illustrative examples of 5-interaction 3-component RD networks showing the five different ways that an external component can be wired into a 2-component RD core, such that the new topology is not an elaboration of one of the minimal architectures in a). Core components and interactions in black, with external in grey. The nature of the net feedback between the external component (BMP in the illustrative examples) and the core positive feedback component (Wnt in the illustrative examples) is shown as a magenta loop for positive feedback and a cyan loop for negative feedback.

c) Examples of 3-component RD networks showing the six different orientations that a set of 3-component RD loops can take, using the 4-interaction minimal architecture in a) 1 as an illustrative example. The feedback loops can be rotated into three different orientations (1, 3 and 5), as shown by the positions of the two feedback loops relative to the dashed black line. In each orientation, the architectures can be reflected around the dashed black line to give a further three orientations (2,4 and 6).

d) Response of Hh to inhibition of Wnt, BMP and Hh for all 45 possible minimal 3-component Wnt-BMP-Hh topologies based on reaction term analysis. Topologies are specified by the combination of a topology (1 to 8 from a) and b)) and an orientation (1 to 6 from c)). Topology architecture 8 is symmetrical upon reflection and therefore two the topology under two possible orientations is the same. The direction of response of Hh shown by arrowhead, with up arrow indicating increase and down arrow decrease. Where the response is unconstrained a white bar is shown. Coupling between components (see supplementary note 4 and fig 4) is not shown. The ten topologies for which the predicted responses are consistent with experimental observations are marked in red. These are the same topologies as recovered by the RD analysis (see fig 3).
