## Supplementary Notes for "Perturbation analysis of a multi-morphogen Turing Reaction-Diffusion stripe patterning system reveals key regulatory interactions"

### 1: Review of Conditions for DDI in 2- 3- and *N*-component systems

The conditions for a diffusion driven instability (DDI) have been extensively analysed. For completeness, we present general conditions for a DDI, as well as specific criteria for assessing whether specific parameterisations of two- and three-component RD systems will give a DDI.

#### *1.1: General conditions for a DDI*

We considered a general reaction-diffusion system of the form

$$\frac{\partial \boldsymbol{u}}{\partial t} = \boldsymbol{f}(\boldsymbol{u}) + \boldsymbol{D}\nabla^2\boldsymbol{u}$$

where $\boldsymbol{u}$ is a vector of $N$ reactant concentrations and $\boldsymbol{D}$ is a diagonal $N$×$N$ matrix of diffusion coefficients where $N \geq 2$, such that

$$\boldsymbol{u} = \begin{pmatrix} u_1 \\ u_2 \\ \vdots \\ u_N \end{pmatrix} \text{ and } \boldsymbol{D} = \begin{pmatrix} D_1 & 0 & \cdots & 0 \\ 0 & D_2 & \cdots & 0 \\ \vdots & \vdots & \ddots & \vdots \\ 0 & 0 & \cdots & D_N \end{pmatrix}.$$

Following the well-established linear stability approach to deriving conditions for DDI, as outlined Murray (2003) (see also White and Gilligan 1998, Marcon et al. 2016), we derive the stability matrix $\boldsymbol{S}$, for which

$$\boldsymbol{S} = \boldsymbol{J} - \boldsymbol{D}q^2,$$

where $q$ is the wave number of a spatially-periodic perturbation, and $\boldsymbol{J}$ is the Jacobian matrix of the reaction system, where the elements $J_{ij}$ are the partial derivatives of the components of $\boldsymbol{f}$, evaluated at a spatially-uniform steady state, such that

$$\boldsymbol{J} = \begin{pmatrix} J_{11} & J_{12} & \cdots & J_{1N} \\ J_{21} & J_{22} & \cdots & J_{2N} \\ \vdots & \vdots & \ddots & \vdots \\ J_{N1} & J_{N2} & \cdots & J_{NN} \end{pmatrix}, \quad J_{ij} = \frac{\partial f_i(u_1,u_2,\ldots,u_N)}{\partial u_j}.$$

Solving the system

$$\det\,[\lambda \boldsymbol{I} - \boldsymbol{S}] = 0$$

gives the dispersion relation for the eigenvalues $\lambda$ of S:

$$\lambda^N + a_1(q^2)\lambda^{N-1} + a_2(q^2)\lambda^{N-2} + \cdots + a_N(q^2) = 0.$$

For diffusion driven instability, the system must be stable without diffusion and unstable with diffusion. Thus, when $q$=0, all solutions $\lambda$ of the dispersion relation must have negative real part, and there must exist positive values of $q^2$ such that there is at least one solution $\lambda$ with positive real part.

We will now consider the specific cases of two- and three-component RD systems.

#### *1.2: Two-component system*

For a two-component system of the form above, the dispersal relationship takes the form

$$\lambda^2 + a_1(q^2)\lambda + a_2(q^2) = 0$$

with coefficients

$$a_1(q^2) = -(J_{11} + J_{22}) + q^2(D_1 + D_2)$$

$$a_2(q^2) = (J_{11}J_{22} - J_{12}J_{21}) - q^2(J_{11}D_2 + J_{22}D_1)$$

Following the approach of Murray (2003) gives the conditions for a DDI that for stability in the absence of diffusion

$$J_{11} + J_{22} < 0$$
$$J_{11}J_{22} - J_{12}J_{21} > 0$$

and for instability with diffusion

$$J_{11}D_2 + J_{22}D_1 > 2\sqrt{D_1D_2(J_{11}J_{22} - J_{12}J_{21})} > 0$$

As is well established (see Murray 2003), if $J_{11}>0$ and $J_{22}<0$, the two components will be in-phase when $J_{12}<0$ and $J_{21}>0$ and out-of-phase when $J_{12}>0$ and $J_{21}<0$.

#### *1.3: Three-component system*

For a three-component system of the form above, the dispersal relationship takes the form

$$\lambda^3 + a_1(q^2)\lambda^2 + a_2(q^2)\lambda + a_3(q^2) = 0$$

with coefficient

$$a_1(q^2) = -(J_{11} + J_{22} + J_{33}) + q^2(D_1 + D_2 + D_3)$$

$$a_2(q^2) = (J_{11}J_{22} - J_{12}J_{21} + J_{11}J_{33} - J_{13}J_{31} + J_{22}J_{33} - J_{23}J_{32}) - q^2(J_1D_2 + J_1D_3 + J_2D_1 + J_2D_3 + J_3D_1 + J_3D_2) + q^4(D_1D_2 + D_1D_3 + D_2D_3)$$

$$a_3(q^2) = -(J_{11}J_{22}J_{33} + J_{12}J_{23}J_{31} + J_{13}J_{32}J_{21} - J_{11}J_{23}J_{32} - J_{22}J_{13}J_{31} - J_{33}J_{12}J_{21}) + q^2((J_{11}J_{22} - J_{12}J_{21})D_3 + (J_{11}J_{33} - J_{13}J_{31})D_2 + (J_{22}J_{33} - J_{23}J_{32})D_1) - q^4(J_{11}D_2D_3 + J_{22}D_1D_3 + J_{33}D_1D_2) + q^6D_1D_2D_3$$

Following the approach of White and Gilligan we considered the Routh-Hurwitz criteria. For stability without diffusion

$$a_1(0) > 0 \wedge a_3(0) > 0 \wedge a_1(0)a_2(0) - a_3(0) > 0$$

and for instability with diffusion

$$a_1(q^2) < 0 \vee a_3(q^2) < 0 \vee a_1(q^2)a_2(q^2) - a_3(q^2) < 0.$$

As detailed by White and Gilligan, the conditions for instability can only be satisfied if

$$a_3(q^2) < 0$$

or

$$a_1(q^2)a_2(q^2) - a_3(q^2) < 0,$$

both of which are cubic equations of the form

$$y(q^2) = a(q^2)^3 + b(q^2)^2 + c(q^2) + d.$$

A DDI will occur if there exists a minimum turning point $q_{TP}^2$, for which $q_{TP}^2 > 0$, and $y(q_{TP}^2) < 0$ (see \White, 1998 #27 for details).

RD systems with more than two components can destabilise either as stable spatially periodic patterns (as for two component systems) or oscillating patterns. When the system is destabilized with real λ the spatial patterns are stationary, while if λ also has imaginary parts, they will oscillate. White and Gilligan provide a detailed analysis of conditions for stable and oscillating patterns. From their analysis, a parameterization will be produce stable spatial patterns for all $q^2$ if

$$a_3(q^2) < 0$$

$$a_1(q^2)a_2(q^2) - a_3(q^2) > 0$$

and

$$a_1(q^2)^2 - a_2(q^2) > 0.$$

The final condition is a quadratic equation in $q^2$ with a minimum at $q^2_{MIN}$, and will be satisfied if $a_1(q^2_{MIN})^2 - a_2(q^2_{MIN}) > 0$.

For each parameterization, a phase relationship was calculated as the eigenvector of the largest positive eigenvalue of the matrix

$$\boldsymbol{S} = \begin{pmatrix} J_{11} - D_1 q^2 & J_{12} & J_{13} \\ J_{21} & J_{22} - D_2 q^2 & J_{23} \\ J_{31} & J_{32} & J_{33} - D_3 q^2 \end{pmatrix}$$

calculated at the turning point $q^2_{TP}$ for which $a_3(q^2) < 0$. While this is not necessarily the phase pattern with which the fastest growing wavelength will destabilise, it does give a phase pattern that is consistent with the network topology defined by the reaction matrix.

#### *1.4: N-component system*

For a general reaction diffusion system, Marcon et al. (2016) demonstrated that coefficients of the dispersal relationship $a_k(q^2)$ for $k$ ={1,…,N} are of the form

$$a_k(q^2) = \sum_{\gamma_k \subseteq S_k^N} \left\{ (-1)^k \det[\boldsymbol{J}(\gamma_k)] + \sum_{m=1}^{k-1} q^{2(k-m)} \sum_{\gamma_m \subset \gamma_k} (-1)^m \det[\boldsymbol{J}(\gamma_m)] \det[\boldsymbol{D}(\tilde{\gamma}_m)] + q^{2k} \det[\boldsymbol{D}(\gamma_k)] \right\} \quad (1.1)$$

where $\gamma_k$ denotes a sequence of k distinct integers $\{i_1, \ldots, i_k\}$, where $1 \leq i_1 < i_2 < \cdots < i_k \leq N$ and $S_k^N$ is the set of all possible sequences of k elements in $\{1, \ldots, N\}$, $\boldsymbol{J}(\gamma_k)$ denotes the $k \times k$ submatrix made up of coefficients with the column and row indices $\gamma_k$, and $\gamma_m$ a sequence of $m < k$ distinct integers, where $\gamma_m \subset \gamma_k$, and $\tilde{\gamma}_m$ is a complementary sequence of integers such that $\gamma_m \cap \tilde{\gamma}_m = 0$ and $\gamma_m \cup \tilde{\gamma}_m = \gamma_k$. As outlined in Marcon et al. (2016), there will be a single real and positive eigenvalue (and stable waves will grow) if $a_k(q^2) > 0$ for $k < N$, and $a_N(q^2) < 0$.

### 2: Perturbation predictions from RD reaction terms

Our analysis of the response of RD systems to perturbations suggested that there is a relationship between the feedback loops within the RD system, and the nature of the response of the system. Specifically, whether a component increases or decreases its level in response to the inhibition of another component depends on whether the components are in a positive or negative feedback loop. As the feedback loops are encoded by the reaction terms of the system, we considered what role the reaction terms could play in this behaviour. Preliminary simulations suggested that the response of an established spatially periodic steady state solution of an RD system to perturbation is predicted by the response of the reaction term equilibrium, and whether it increases or decreases in response to equivalent perturbations (data not shown). We therefore decided to further investigate what governed the responses of the reaction terms.

#### *2.1: Shape of N-component response*

We considered the response of a system of $N$ components, using a system of linear ODEs equivalent to that used for the full reaction-diffusion system we used before, where

$$\frac{du_i}{dt} = \sum_{j=1}^{N} a_{ij}u_j + b_i - c_iu_i.$$

The parameters $a_{ij}$ represent the weight of the interaction from component $u_j$ to component $u_i$, where a positive term indicates an activation and a negative term an inhibition, $b_i$ is a background production term and $c_i$ is the linear degradation rate of component $u_i$.

The position of the equilibrium is given by setting $\frac{du_i}{dt} = 0$ for all $i$, and as in (Desoer 1960), rearranging this gives

$$u_i = (\boldsymbol{A}^{-1}\boldsymbol{b})_i = -\frac{\sum_{j=1}^{N} \mathrm{C}[\boldsymbol{A}]_{ji} b_j}{\det[\boldsymbol{A}]}$$

where matrix $\boldsymbol{A}$ given by

$$\boldsymbol{A} = \begin{pmatrix} a_{11} - c_1 & a_{12} & \cdots & a_{1N} \\ a_{21} & a_{22} - c_2 & \cdots & a_{2N} \\ \vdots & \vdots & \ddots & \vdots \\ a_{N1} & a_{N2} & \cdots & a_{NN} - c_N \end{pmatrix}$$

and $\mathrm{C}[\boldsymbol{A}]_{ji}$ is the cofactor of the element of the $j^{\text{th}}$ row and $i^{\text{th}}$ column of $\boldsymbol{A}$.

We considered the effect of inhibiting a component on the equilibrium levels of the system. Specifically, we looked at the effect of inhibiting component $u_1$ on both the levels of $u_1$ itself, and the levels of other components by looking at the effect on a component $u_2$. As for our simulations of full RD systems we performed two forms of inhibition, namely inhibiting the response of a component to the levels of $u_1$ (analogous to inhibiting a "$u_1$ receptor") and inhibiting the production of $u_1$.

2.1.1: Inhibition of response to $u_1$

We first consider the effect of perturbing the response to $u_1$ by reducing the weight of all interactions from component $u_1$ (to itself and all other components) by a factor of $1 - \alpha$ (ie with no inhibition when $\alpha = 0$ and a full inhibition when $\alpha = 1$). At equilibrium, this gave a system of the form

$$\frac{du_1}{dt} = 0 = (1 - \alpha)a_{11}u_1 + \sum_{j=2}^{N} a_{1j}u_j + b_1 - c_1u_1$$

and

$$\frac{du_i}{dt} = 0 = (1 - \alpha)a_{i1}u_i + \sum_{j=2}^{N} a_{ij}u_j + b_i - c_iu_i,$$

for $i$={2,…,$N$}. It should be noted that for the response of $u_1$ to its own inhibition, only the weight of $a_{11}$ is reduced, as $c_1$ represents a linear degradation term and would therefore be unaffected directly by an inhibitor blocking a receptor.

Rearranging these equations gave equilibrium levels of $u_i$ upon inhibition $\alpha$, where

$$u_1 = -\frac{\sum_{j=1}^{N} \mathrm{C}[\boldsymbol{A}]_{j1} b_j}{\det[\boldsymbol{A}] - \alpha \det[\boldsymbol{C}]} \quad (2.1)$$

and

$$u_i = -\frac{\sum_{j=1}^{N} \mathrm{C}[\boldsymbol{A}]_{ji} b_j - \alpha \sum_{j=1}^{N} \mathrm{C}[\boldsymbol{C}]_{ji} b_j}{\det[\boldsymbol{A}] - \alpha \det[\boldsymbol{C}]} \quad (2.2)$$

for $i$={2,...,N}. Matrix **C**, which does not include the degradation term $c_1$, is given by

$$\boldsymbol{C} = \begin{pmatrix} a_{11} & a_{12} & \cdots & a_{1N} \\ a_{21} & a_{22} - c_2 & \cdots & a_{2N} \\ \vdots & \vdots & \ddots & \vdots \\ a_{N1} & a_{N2} & \cdots & a_{NN} - c_N \end{pmatrix}$$

and $\mathrm{C}[\boldsymbol{C}]_{ji}$ is the cofactor of the element of the jth row and ith column of $\boldsymbol{C}$.

*Response of $u_1$:*

The response of $u_1$ to its own inhibition is governed by equation 2.1, which is a function of $\alpha$ with horizontal and vertical asymptotes at

$u_1 = 0$

and

$\alpha = \frac{\det[\boldsymbol{A}]}{\det[\boldsymbol{C}]}$,

respectively. The intercept of the $u_1$-axis lies at

$$u_1 = -\frac{\sum_{j=1}^{N} \mathrm{C}[\boldsymbol{C}]_{j1} b_j}{\det[\boldsymbol{A}]},$$

which is the unperturbed level of $u_1$ (i.e. $\alpha = 0$). As the unperturbed level of $u_1$ has a positive value, the function can take two forms depending on whether the vertical asymptote takes a positive or negative value (see figure S1 a).

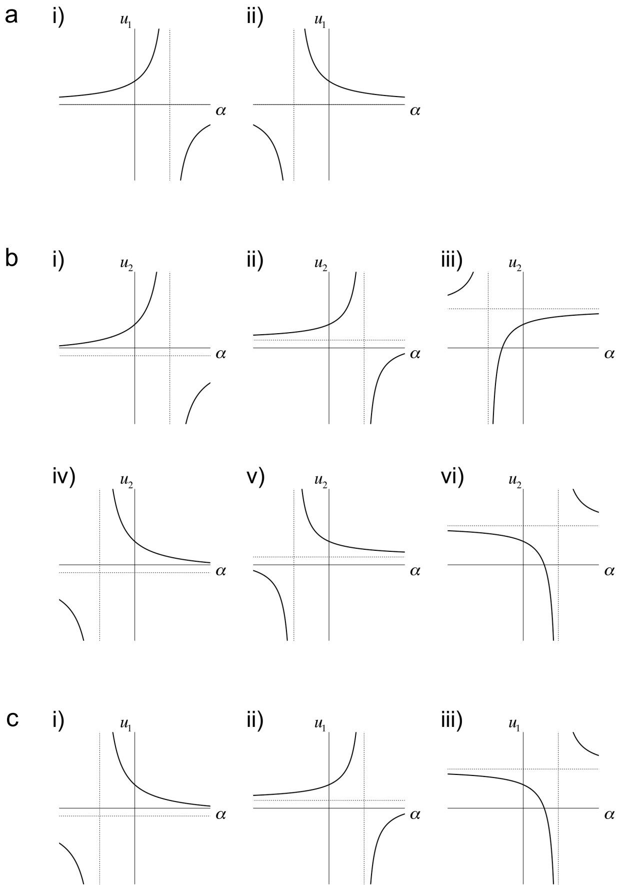


Figure S1
a) Graphs showing the two possible responses of $u_1$ to the inhibition of $u_1$ by strength $\alpha$. The vertical asymptote (dashed line) can lie at a positive (i) or negative (ii) value of $\alpha$. The horizontal asymptote lies along the $\alpha$-axis.
b) Graphs showing the six possible responses of $u_2$ to the inhibition of $u_1$ by strength $\alpha$. The vertical asymptote (vertical dashed line) can lie at a positive (i, ii and vi) or negative (iii, iv and v) value of $\alpha$. The horizontal asymptote (horizontal dashed line) can lie at a positive (ii, iii, v and vi) or negative (i and iv) value of $u_2$. The intercept of the $\alpha$-axis can lie at a positive (ii, iv and vi) or negative (i, iii and v) value of $\alpha$.
c) Graphs showing the three possible responses of $u_1$to the inhibition of $u_1$ by strength $\alpha$. The vertical asymptote (vertical dashed line) can lie at a positive (ii and iii) or negative (i) value of $\alpha$. The horizontal asymptote (horizontal dashed line) can lie at a positive (ii and iii) or negative (i) value of $u_1$. The intercept of the $\alpha$-axis can lies at $\alpha$=1.

Therefore, $u_1$ will increase with its own inhibition if

$$\frac{\det[\boldsymbol{A}]}{\det[\boldsymbol{C}]} > 0 \quad (2.3)$$

and will decrease if

$$\frac{\det[\boldsymbol{A}]}{\det[\boldsymbol{C}]} < 0. \quad (2.4)$$

*Response of $u_2$:*

Likewise, the response of $u_2$ to inhibition of $u_1$ is governed by equation 2.2, which is also a function of $\alpha$ with horizontal and vertical asymptotes, which lie at

$$u_2 = -\frac{\sum_{j=1}^{N} \mathrm{C}[\boldsymbol{C}]_{j2} b_j}{\det[\boldsymbol{C}]}$$

and

$$\alpha = \frac{\det[\boldsymbol{A}]}{\det[\boldsymbol{C}]},$$

respectively. The intercept of the $u_2$-axis lies at

$$u_2 = -\frac{\sum_{j=1}^{N} \mathrm{C}[\boldsymbol{A}]_{j2} b_j}{\det[\boldsymbol{A}]},$$

which is the unperturbed state and has a positive value, while the intercept of the $\alpha$-axis is at

$$\alpha = \frac{\sum_{j=1}^{N} \mathrm{C}[\boldsymbol{A}]_{ji} b_j}{\sum_{j=1}^{N} \mathrm{C}[\boldsymbol{C}]_{ji} b_j}.$$

Depending on the relative values of the asymptotes and intercepts, this function can take six forms (figure S1 b), for three of which (figure S1 b i-iii), $u_2$ increases with inhibition of $u_1$, while for the remaining three (figure S1 b iv-vi), $u_2$ decreases with the inhibition of $u_1$. From the relative positions of the horizontal asymptote relative to the unperturbed state at $\alpha = 0$, for conditions i, ii, iv and v

$$-\frac{\sum_{j=1}^{N} \mathrm{C}[\boldsymbol{A}]_{j2} b_j}{\det[\boldsymbol{A}]} > -\frac{\sum_{j=1}^{N} \mathrm{C}[\boldsymbol{C}]_{j2} b_j}{\det[\boldsymbol{C}]},$$

while for conditions iii and vi

$$-\frac{\sum_{j=1}^{N} \mathrm{C}[\boldsymbol{A}]_{j2} b_j}{\det[\boldsymbol{A}]} < -\frac{\sum_{j=1}^{N} \mathrm{C}[\boldsymbol{C}]_{j2} b_j}{\det[\boldsymbol{C}]}.$$

However, from the position of the vertical asymptote, for i, ii and vi

$$\frac{\det[\boldsymbol{A}]}{\det[\boldsymbol{C}]} > 0,$$

while for iii, iv and v

$$\frac{\det[\boldsymbol{A}]}{\det[\boldsymbol{C}]} < 0.$$

Therefore, in conditions i, ii and iii where $u_2$ increases on the inhibition of $u_1$

$$\det[\boldsymbol{C}] \sum_{j=1}^{N} \mathrm{C}[\boldsymbol{A}]_{j2} b_j < \det[\boldsymbol{A}] \sum_{j=1}^{N} \mathrm{C}[\boldsymbol{C}]_{j2} b_j \quad (2.5)$$

while in conditions iv, v and vi where $u_2$ decreases on the inhibition of $u_1$

$$\det[\boldsymbol{C}] \sum_{j=1}^{N} \mathrm{C}[\boldsymbol{A}]_{j2} b_j > \det[\boldsymbol{A}] \sum_{j=1}^{N} \mathrm{C}[\boldsymbol{C}]_{j2} b_j. \quad (2.6)$$

#### 2.1.2: Inhibition of production of $u_1$

We next considered the effect of inhibiting the production of $u_1$ by reducing the weights of all the coefficients effecting the production of $u_1$ (including the constant term, but excluding the degradation coefficient) by a factor of 1-$\alpha$. At equilibrium, this gives a system of the form

$$\frac{du_1}{dt} = 0 = (1-\alpha)(\sum_{j=1}^{N} a_{1j}u_j + b_1) - c_1u_1$$

and

$$\frac{du_i}{dt} = 0 = \sum_{j=1}^{N} a_{ij}u_j + b_i - c_iu_i$$

for $i$={2,...,$N$}. Rearranging these equations gives the equilibrium levels of $u_i$ upon inhibition $\alpha$, where

$$u_1 = -\frac{\sum_{j=1}^{N} \mathrm{C}[\boldsymbol{A}]_{j1} b_j (1-\alpha)}{\det[\boldsymbol{A}] - \alpha \det[\boldsymbol{C}]} \qquad (2.9)$$

and

$$u_i = -\frac{\sum_{j=1}^{N} \mathrm{C}[\boldsymbol{A}]_{ji} b_j - \alpha \sum_{j=1}^{N} \mathrm{C}[\boldsymbol{C}]_{ji} b_j}{\det[\boldsymbol{A}] - \alpha \det[\boldsymbol{C}]}$$

for $i$={2,...,$N$}. As this equation describing the behaviour of $u_i$ where $i \neq 1$ is the same as for the inhibition of response (equation 2.2), the shape of the response of $u_2$ to an inhibition of strength a will be the same for inhibiting the production of or response to $u_1$.

*Behaviour of* $u_1$*:*

The behaviour of $u_1$ is again governed by a function of $\alpha$ (equation 2.9) with horizontal and vertical asymptotes, which now lie at

$$u_1 = -\frac{\sum_{j=1}^{N} \mathrm{C}[\boldsymbol{A}]_{j1} b_j}{\det[\boldsymbol{C}]}$$

and

$$\alpha = \frac{\det[\boldsymbol{A}]}{\det[\boldsymbol{C}]},$$

respectively. The intercept of the $u_1$-axis lies at

$$u_1 = -\frac{\sum_{j=1}^{N} \mathrm{C}[\boldsymbol{A}]_{j1} b_j}{\det[\boldsymbol{A}]}$$

which is the unperturbed state and has a positive value, while the intercept of the $\alpha$-axis is at

$$\alpha = 1.$$

Depending on the relative values of the asymptotes and intercepts, this function can take three forms (figure S1 c), for two of which (S1 c i, iii), $u_1$ decrease with its inhibition, while for the remaining one (figure S1 c ii), $u_1$ increases with its inhibition. Therefore, $u_1$ will increase on its inhibition if

$$0 > \frac{\det[\boldsymbol{A}]}{\det[\boldsymbol{C}]} \quad (2.10)$$

or

$$1 < \frac{\det[\boldsymbol{A}]}{\det[\boldsymbol{C}]}. \quad (2.11)$$

While $u_1$ will decrease on its inhibition if

$$0 < \frac{\det[\boldsymbol{A}]}{\det[\boldsymbol{C}]} < 1. \quad (2.12)$$

#### *2.2: Comparison with a numerically modelled RD system*

Using this framework, we could compare the responses to inhibition for the 2- and 3-component RD systems with the behaviour of the reaction terms. We took the parameterisations of the reaction terms used in the full RD simulations (fig 3 and fig 3 supplement 2) and inferred the direction of response by calculating the relative position of the asymptotes and intercepts. Comparing the direction of the response to an inhibition for the reaction terms to the full RD system (figure S2) showed that the pattern of the responses is very similar. This suggests that the response to an inhibition is predominantly dictated by the response of the reaction terms.

What about the small proportion of parameterisations behave differently between the full RD system and the reaction terms alone? Interestingly, repeating the simulations, but reducing the strength of the inhibition by a factor of 0.25 or $0.25^2$, successively increased the discrepancy (figure S2). As decreasing the strength of an inhibition decreases the magnitude of the response for the reaction terms, this suggests that the response of the reaction terms will dictate the response of the RD system provided that they are sufficiently large. As the only difference between the reaction term systems and the full RD systems is diffusion, this suggests that if the response to an inhibition of the reaction terms is not sufficiently large, there will be a significant effect from diffusion, which in some cases can change the direction of response.

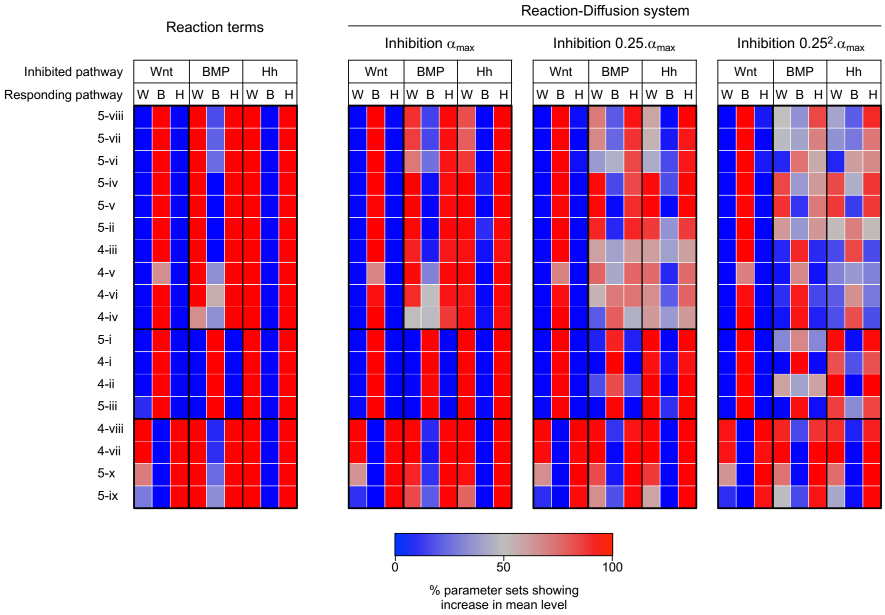


Figure S2
Full maps showing the percentage of parameter sets where the level of each component increases in response to the inhibition of each component in the network in reaction term analysis and RD simulations. Left hand heat map shows response for reaction terms analysis of the parameter sets used in 3-component RD analysis, evaluated using criteria from supplementary note 2. Heat map showing Reaction-Diffusion system with inhibition strength $\alpha_{max}$ is a reproduction of the heat map from figure 3, and heat maps with inhibition strength $0.25.\alpha_{max}$ and $0.25^2.\alpha_{max}$ are from reruns of the 3-component simulations replacing the inhibition parameter value $\alpha$ with values of $0.25.\alpha$ and $0.25^2.\alpha$ respectively. Topology names are as in figure 3.

This is likely to be generally true for RD systems based on the following argument. Large perturbations in reaction terms will produce large shifts in the mean value of the periodic pattern while changes in the diffusion terms will flatten or sharpen the peaks and troughs. For example, an increase in the reaction term will increase the value of $u_i$ in both peaks and troughs. For equilibrium to be restored, the diffusion term $D_i \frac{\partial^2 u_i}{\partial x^2}$ has to decrease. Since at the peaks, $D_i \frac{\partial^2 u_i}{\partial x^2}$ is negative, this means it has to become more negative, i.e. the peaks must become sharper. Conversely, the troughs must become flatter (second derivative is positive becomes less positive). These changes in the curvature of the peaks and troughs do not necessarily have a net effect on the average levels that are the same or opposite to the reaction term change. Our simulations, however, say that their effects do not increase at the same rate as those of the reaction terms. The simulations suggest that their role is more pronounced at small reaction term changes although we have not been able to demonstrate this analytically.

### 3: A graphical interpretation of the responses to perturbation

We wanted to understand how the position of a component in a network affects the responses of the system to its inhibition, in particular, relative to the type of feedback loop the component is found in. In a recent study Marcon et al. (2016) used such a graphical approach to understand the conditions for DDI, in terms of stabilizing and destabilizing contributions of different feedback loops. This was based on an interpretation of the determinant as a combination of the cycles defined by the reaction matrix. As the conditions determining the response to perturbation (see supplementary note 2) are in part dependent on the determinants of matrices $\boldsymbol{A}$ and $\boldsymbol{C}$, we took a similar approach.

In brief, a matrix $\boldsymbol{M}$ (such as the reaction matrices $\boldsymbol{A}$ and $\boldsymbol{C}$ or in the case of Marcon et al. the Jacobian matrix $\boldsymbol{J}$) has associated graph Gr[$\boldsymbol{M}$] which is a labelled, weighted, directed graph of $N$ nodes with edges from node $j$ to node $i$ of weight $a_{ij}$ where $a_{ij} \neq 0$. This is the graphical representation of the reaction network which we have been using throughout to depict of the signs of the interactions as a network topology. Such a graph contains cycles, where a path can be traced from any node, through a series of other nodes, back to itself (or directly onto itself without passing through any other nodes). A cycle represents a positive feedback loop if the product of the weights of the edges is positive, and a negative feedback loop if the product is negative.

For any graph such as Gr[$\boldsymbol{M}$], a subgraph can be defined containing all nodes but only a subset of edges where each node receives an input from only one other node and also outputs to only one other node (i.e. each node has both an indegree and outdegree of 1). Such a subgraph is called a linear spanning subgraph (L-subgraph, $l$), and is made up of a set of 1 to $N$ cycles through all nodes. The weight of an L-subgraph $w(l)$ is given by the product of the weights of these cycles, with the weight of each cycle itself the product of the edges within. The determinant of the $N \times N$ matrix $\boldsymbol{M}$ is given by the sum of the weights of each L-subgraph $l$, in graph Gr[$\boldsymbol{M}$], with signs as below

$$\det[\boldsymbol{M}] = (-1)^N \sum_{l \subseteq Gr} (-1)^{\sigma(l)} w(l), \quad (3.1)$$

where $\sigma(l)$ is the number of cycles in a L-spanning subgraph. This is known as the Coates formula for more details on the derivation see Marcon et al. (2016) and references therein.

#### *3.1: Response of $u_1$*

As demonstrated in supplemental note 2, the response of $u_1$ to its own inhibition depends on the relative signs and magnitudes of the determinant det[$\boldsymbol{C}$] relative to det[$\boldsymbol{A}$], the sign of which itself is determined by the criteria for a DDI (see supplementary note 1). To understand the response of $u_1$ to its own inhibition in graphical terms, we therefore considered these determinants in terms of the feedback loops (or cycles) through $u_1$ in matrix $\boldsymbol{C}$. From equation 3.1 each summand in det [$\boldsymbol{C}$] can be written in the form $(-1)^N(-1)^{\sigma(l)}w(l)$, where $l$ is a linear spanning subgraph of Gr[$\boldsymbol{C}$]. As all nodes in an $l$-graph have an indegree and an outdegree of 1, for each summand in equation 3.1 there is a single cycle containing node $u_1$, which comprises a set of $n$ nodes, where $1 \leq n \leq N$, with $\kappa_n$ a sequence of integers $\{i_1,i_2,\ldots,i_n\}$ which defines the series which the cycles passes through the nodes $u_i$, with each such sequence starting at $i_1$=1. If $n$=1, the cycle forms an interaction from $u_1$ directly onto itself, while if $n=N$, the cycle passes through all nodes. $S_{\kappa_n}$ is the set of all possible $\kappa_n$, and $w(\kappa_n)$ is the weight of the cycle defined by $\kappa_n$ (positive for a positive feedback loop, negative for a negative feedback loop). For each such cycle, where $n<N$, the nodes excluded from the cycle form as L-graph $l_n$ consisting of 1 to $N$-$n$ cycles between the remaining nodes, where $\sigma(l_n)$ is the number of remaining cycles (with $\sigma(l_n) = \sigma(l) - 1$ as one cycle has been remove). $w(l_n)$ is the product of the weights of the remaining cycles such that $w(l) = w(\kappa_n)w(l_n)$, and $S_{l_n}$ is the set of all possible $l$-graphs $l_n$ for a given $\kappa_n$.

Therefore, for any term in the determinant, the loop through $u_1$ can be factored out to give

$$(-1)^N(-1)^{\sigma(l)}w(l) = -(-1)^n w(\kappa_n)(-1)^{N-n}(-1)^{\sigma(l_n)}w(l_n).$$

Gathering together determinant terms containing the same cycle through $u_1$, the determinant can be rewritten

$$\det[\boldsymbol{C}] = -\sum_{\kappa_n \subseteq S_{\kappa_n}} (-1)^n w(\kappa_n)(-1)^{N-n} \sum_{l_n \subseteq S_{l_n}} (-1)^{\sigma(l_n)} w(l_n)$$

$$= -\sum_{\kappa_n \subseteq S_{\kappa_n}} (-1)^n w(\kappa_n) \det[\boldsymbol{C}(\tilde{\kappa}_n)], \quad (3.2)$$

where $\boldsymbol{C}(\tilde{\kappa}_n)$ is the submatrix of $\boldsymbol{C}$ excluding the columns and rows defined in $\kappa_n$ (for n=N, $\det[\boldsymbol{C}(\tilde{\kappa}_n)] = 1$ as it is the determinant of an 0x0 matrix).

3.1.1: Inhibiting response to $u_1$

As demonstrated in supplementary note 2, the behaviour of $u_1$ on inhibiting the response to $u_1$ is dependent on whether the sign of $\frac{\det[\boldsymbol{A}]}{\det[\boldsymbol{C}]}$ is positive or negative (see inequalities 2.3 and 2.4). Therefore, $u_1$ will increase on its own inhibition if

$$\frac{\sum_{\kappa_n \subseteq S_{\kappa_n}} (-1)^n w(\kappa_n) \det[\boldsymbol{C}(\tilde{\kappa}_n)]}{\det[\boldsymbol{A}]} < 0 \quad (3.3)$$

and will decrease if

$$\frac{\sum_{\kappa_n \subseteq S_{\kappa_n}} (-1)^n w(\kappa_n) \det[\boldsymbol{C}(\tilde{\kappa}_n)]}{\det[\boldsymbol{A}]} > 0, \quad (3.4)$$

where each summand contains a different cycle through $u_1$, each multiplied by the determinant of the submatrix made of component excluded from the cycle. The response of $u_1$ to the inhibition of its response therefore depends on the natures of the feedback loops through $u_1$, and the stability of the components excluded from the loops.

#### 3.1.2: Inhibiting production of $u_1$

For inhibiting the production of $u_1$, we demonstrated in supplementary note 2 that the response of $u_1$ again depends on det $[\boldsymbol{A}]$ and det $[\boldsymbol{C}]$ (see inequalities 2.10 – 2.12). First considering when $N$ is even, from supplementary note 1 it follows that det $[\boldsymbol{A}] > 0$. To satisfy the condition 2.10, det $[\boldsymbol{C}] < 0$, and as a consequence, det $[\boldsymbol{A}] >$ det $[\boldsymbol{C}]$. Otherwise, det $[\boldsymbol{C}] > 0$, and in this case, to also satisfy condition 2.11, it follows that again det $[\boldsymbol{A}] >$ det $[\boldsymbol{C}]$. However, when condition 2.12 holds, it follows that det $[\boldsymbol{A}] <$ det $[\boldsymbol{C}]$. Therefore, when det $[\boldsymbol{A}] <$ det $[\boldsymbol{C}]$, $u_1$ increases when its production is inhibited, while when det$[\boldsymbol{A}] >$ det $[\boldsymbol{C}]$, $u_1$ decreases when its production is inhibited.

As matrices $\boldsymbol{A}$ and $\boldsymbol{C}$ only differ in the presence of absence of the $-c_1$ in the first matrix term, then

$$\det[\boldsymbol{A}] = \det[\boldsymbol{C}] - c_1 \det[\boldsymbol{C}(\tilde{\kappa}_1)],$$

where $\boldsymbol{C}(\tilde{\kappa}_1)$ is the submatrix of $\boldsymbol{C}$, excluding the first row and column. If therefore follows that $u_1$ will increase on its inhibition if

$$c_1 \det[\boldsymbol{C}(\tilde{\kappa}_1)] > 0,$$

and $u_1$ will decrease on its inhibition if

$$c_1 \det[\boldsymbol{C}(\tilde{\kappa}_1)] < 0.$$

Given that $c_1 > 0$, it follows that $u_1$ will increase when its production is inhibited if

$$\frac{\det[\boldsymbol{C}(\tilde{\kappa}_1)]}{\det\,[\boldsymbol{A}]} > 0 \quad (3.5)$$

and will decrease when

$$\frac{\det[\boldsymbol{C}(\tilde{\kappa}_1)]}{\det\,[\boldsymbol{A}]} < 0. \quad (3.6)$$

By the same reasoning, these conditions also hold if $N$ is odd. The response of $u_1$ to the inhibition of its production therefore depends on the stability of the submatrix made from the remaining components.

***3.2: Response of $u_2$:***

As demonstrated in supplementary note 2 (conditions 2.5 and 2.6), the responses of component $u_2$ in an N-component system to the inhibition of $u_1$ is in part determined by the relative signs and sizes of the determinants det[$\boldsymbol{A}$] and det[$\boldsymbol{C}$]. The responses to inhibition also depend on the terms, $\sum_{j=1}^{N} \mathrm{C}[\boldsymbol{A}]_{j2} b_j$ and $\sum_{j=1}^{N} \mathrm{C}[\boldsymbol{C}]_{j2} b_j$ which are of the same form as a determinant, but with the second column of matrix $\boldsymbol{A}$ and $\boldsymbol{C}$ respectively replaced by the vector of background production terms ($b_1$,$b_2$,...,$b_N$). Therefore, each term in these sums can similarly be interpreted graphically, as a path from an external node $u_b$ to component $u_2$, with all components not involved in the path making up a set of cycles with each node having an indegree and an outdegree of 1. Therefore, for each summand, every node has a single input and a single output, except $u_b$ which only has a single output and no input, and $u_2$ which only has a single input and no output.

These conditions 2.5 and 2.6 can be rearranged to give

$$c_1 \sum_{j=2}^{N} \mathrm{C}[\boldsymbol{C}(\tilde{\kappa}_1)]_{(j-1)1} a_{j1} \sum_{j=1}^{N} \mathrm{C}[\boldsymbol{A}]_{j1} b_j > 0 \quad (3.7)$$

and

$$c_1 \sum_{j=2}^{N} \mathrm{C}[\boldsymbol{C}(\tilde{\kappa}_1)]_{(j-1)1} a_{j1} \sum_{j=1}^{N} \mathrm{C}[\boldsymbol{A}]_{j1} b_j < 0, \quad (3.8)$$

respectively, where $\mathrm{C}[\boldsymbol{C}(\tilde{\kappa}_1)]_{(j-1)1}$ is the cofactor of the term $a_{j2}$ from matrix $\boldsymbol{C}$ in the submatrix $\boldsymbol{C}(\tilde{\kappa}_1)$ (as this submatrix excludes the first row and column of $\boldsymbol{C}$, this will be the cofactor of the first column and the ($j$-1)th row).

The only term in conditions 3.7 and 3.8 whose sign is not constrained is $\sum_{j=2}^{N} \mathrm{C}[\boldsymbol{C}(\tilde{\kappa}_1)]_{(j-1)1} a_{j1}$. From equations 2.1 and 2.9 which describe the levels of $u_1$ under inhibition, for the unperturbed state of $u_1$ to be positive, the term $\sum_{j=1}^{N} \mathrm{C}[\boldsymbol{A}]_{j1} b_j$ must have the same sign as the determinant $\det[\boldsymbol{A}]$ (positive if there are an even number of components and negative if there are an odd number). Also, $-c_1 < 0$ as it represents a degradation term. The form of the sum $\sum_{j=2}^{N} \mathrm{C}[\boldsymbol{C}(\tilde{\kappa}_1)]_{(j-1)1} a_{j1}$ is the same as that of $\sum_{j=1}^{N} \mathrm{C}[\boldsymbol{A}]_{j1} b_j$, and so it can be interpreted as a sum where each term represents a path from component $u_1$ to component $u_2$. Specifically,

$$\sum_{j=2}^{N} \mathrm{C}[\boldsymbol{A}(\tilde{\kappa}_1)]_{(j-1)1} a_{j1} = -\sum_{p_n \subseteq S_{p_n}} (-1)^n w(p_n) \det[\boldsymbol{A}(\tilde{\kappa}_n)], \quad (3.9)$$

where $p_n$ is a sequence of integers $\{i_1, i_2, \ldots, i_n\}$, where $n \leq 2 \leq N$, starting with $i_1$=1 and ending with $i_n$=2, which defines the series of nodes $u_i$ through which the path from $u_1$ to $u_2$ passes, $S_{p_n}$ the set of all possible such paths, and with $w(p_n)$ the weight of the path. The response of $u_2$ to the inhibition of $u_1$ therefore depends on the weight of the paths between the two nodes, and the stability of the components excluded from the paths.

**Supplementary note 4: Constraints on minimal topologies**

Having established graphical conditions for the response of a system to inhibition of each component, we wanted to establish how the response of the system is related to network topology. As with our numerical investigation of 3-component systems, we set out to understand the behavior of the subset of networks with the minimal requirements for patterning. We first establish the minimal requirements for a strongly-connected RD system, before considering how these networks respond to perturbation.

***4.1: Identifying minimal topologies***

<u>4.1.1: Minimal conditions for a DDI</u>

In order to establish the minimal requirements for RD we asked what are the fewest interactions (excluding degradation terms) and feedback loops needed to satisfy the criteria for DDI. It should be noted that as we have been using a system of linear equations in our Reaction-Diffusion system, the reaction matrix $\boldsymbol{A}$ is the same as the Jacobian matrix $\boldsymbol{J}$, and the number of interactions is given by the number of nonzero terms $a_{ij}$. As described in Supplementary note 1, conditions for DDI that will always give rise to stable waves require that $a_N(q^2) < 0$. Considering the terms that make up $a_N(q^2)$ (equation 1.1), the first term is made up solely of reaction terms and requires that $(-1)^N\det[\boldsymbol{J}]>0$ in order to satisfy the criterion of stability in the absence of diffusion ($a_N(0) > 0$). The third term is made solely of diffusion coefficients and the wave length coefficient $q^2$, both of which are positive, meaning that $q^{2N}det[\boldsymbol{D}] > 0$. For the remaining terms, which contain both reaction and diffusion coefficients, as above for the diffusion and wavelength coefficients $q^{2(N-m)}\det[\boldsymbol{D}(\bar{\gamma}_m)] > 0$. Therefore, the conditions for DDI depend on the reaction terms and the signs of the various terms $(-1)^m\det[\boldsymbol{J}(\gamma_m)]$.

We first considered a system consisting of $N$ components, without any interactions between them. These components will be only undergoing degradation (i.e. the only nonzero terms in $\boldsymbol{J}$ are on the diagonal, and are negative or in terms of reaction matrix $\boldsymbol{A}$, for all coefficients $a_{ij}$=0). From equation 3.1, the only nonzero summands of any of the submatrix determinant $\det[\boldsymbol{J}(\gamma_m)]$ in equation 1.1, will be those consisting only self-interactions (i.e. the degradation terms). It therefore follows that

$$(-1)^m \det[\boldsymbol{J}(\gamma_m)] = (-1)^m \prod_{i=1}^{m} J_{(\gamma_m)_i(\gamma_m)_i} \quad (4.1)$$

which will always have a positive sign. Therefore, unsurprisingly, the condition for instability when there is diffusion, as $a_N(q^2) < 0$, cannot be satisfied in this simple system.

In order to satisfy the conditions for DDI there must be at least one term in $a_N(q^2)$ with a negative sign. From equation 4.1 this can be achieved by simply adding a single autoactivation term for a component. For example, if $u_1$ directly activates itself, providing that this autoactivation is strong enough to outweigh the degradation term for $u_1$ (in terms of matrix $\boldsymbol{A}$, $a_{11} - c_1 > 0$) a single term on the diagonal of the Jacobian matrix $J_{11}$ will go from being negative to positive, resulting in the sign of any term $det[\boldsymbol{J}(\gamma_m)]$ for which $\gamma_m$ contains $u_1$ also changing sign. Alternatively, this can also be achieved by the addition of a single positive feedback loop through the set of $r$ nodes defined by the sequence $\gamma_r$ of weight $w(\gamma_r)$. For any $\det[\boldsymbol{J}(\gamma_m)]$ where $\gamma_r$ is a subset of $\gamma_m$

$$\det[\boldsymbol{J}(\gamma_m)] = \left(\prod_{i=1}^{r} J_{(\gamma_r)_i(\gamma_r)_i} - (-1)^r w(\gamma_r)\right) \prod_{j=1}^{m-r} J_{(\tilde{\gamma}_r)_j(\tilde{\gamma}_r)_j}$$

where $\tilde{\gamma}_r$ is a complementary sequence of integers to $\gamma_r$ such that $\gamma_r \cap \tilde{\gamma}_r = 0$ and $\gamma_r \cup \tilde{\gamma}_r = \gamma_m$. Again, provided that the weight of the positive feedback loop is greater than for the degradation terms, $\det[\boldsymbol{J}(\gamma_m)]$ will change sign.

However, as the sign of any submatrix containing the positive feedback loop will change sign, it follows that with the addition of a single destabilising positive feedback loop, $(-1)^N \det[\boldsymbol{J}]<0$. This will no longer satisfy the condition for stability in the absence of diffusion $a_N(0) > 0$. An additional positive summand in $a_N(0)$ is therefore needed to stabilize the system. By the above reasoning, this can most simply be achieved by the addition of a single negative feedback between a set of components $\gamma_s$, where $\gamma_s \neq \gamma_r$. If this negative feedback loop takes the form of a path from a node $\gamma_r$, through the remaining nodes in the system and back into any node in $\gamma_r$, a single additional term will be added to any $\det[\boldsymbol{J}(\gamma_m)]$ where $\gamma_s$ is a subset of $\gamma_m$ (including those for which $\gamma_r$ is also a different subset). If the weight of this loop is sufficiently large, then it overcome the effect of the positive feedback loop, and satisfy $a_N(0) > 0$.

Moreover, as there will still exist $\gamma_m$ for which $\gamma_r$ is a subset, but not $\gamma_s$, there will be terms $q^{2(k-m)}(-1)^m \det[\boldsymbol{J}(\gamma_m)]\ \det[\boldsymbol{D}(\tilde{\gamma}_m)]$ which will remain negative, even after the addition of a negative feedback loop to the system. Therefore, with the appropriate weighting of diffusion coefficients, there will exist conditions where $a_k(q^2) < 0$ when $k$=$N$, but not for other $k$, while maintaining $a_N(0) > 0$. As such $\tilde{\gamma}_m$ will by definition exclude components of the positive feedback loop through the nodes in $\gamma_r$, to achieve such a weighing, diffusion coefficients among components in the negative will be larger than those in the positive feedback loop. This gives an intuitive understanding for then general idea of local autoactivation and long-range inhibition in RD systems. In summary, a system containing single positive feedback loop coupled to a negative feedback loop is, therefore, sufficient to satisfy the conditions for a DDI for any size system, given appropriate strengths of interactions and diffusion coefficients.

Considering the minimal requirements for DDI in terms of the number of interactions between components in interaction matrix $\boldsymbol{A}$ (including autoactivation terms $a_{ii}$, but not degradation terms $c_i$), in any such system, a positive feedback loop through $r$ components will add $r$ interactions to the system, while the negative feedback loop though the remaining $N$-$r$ components will add a further $N$-$r$+1 terms, resulting in a total of $N$+1 interactions. For any strongly-connected network, it is clear than fewer than $N$+1 interactions will not allow sufficient feedback loops to be made to satisfy the criteria for DDI, while other networks made of $N$+1 interactions (e.g. two positive or two negative feedback loops) cannot satisfy the conditions for DDI. While there are other ways of satisfying these conditions, they will require additional feedback loops, and therefore additional interaction. Therefore, in a system of $N$ components, the combination of a single positive feedback loop through one to $N$-1 components, and a negative feedback loop through the remaining components is the minimal requirement for a DDI.

4.1.2: The phase of minimal DDI systems

Having established the architecture of minimal topologies, we next sought to establish the relative phase of the components for each of these topologies. This is dictated by the signs of the eigenvector of the largest positive eigenvalue of the Jacobian matrix – components with the same sign will be in the same phase. From the derivation of conditions for DDI in supplementary note 1, we have the relationship

$$\begin{pmatrix} J_{11} - D_1 q^2 - \lambda & J_{12} & \cdots & J_{1N} \\ J_{21} & J_{22} - D_2 q^2 - \lambda & \cdots & J_{2N} \\ \vdots & \vdots & \ddots & \vdots \\ J_{N1} & J_{N2} & \cdots & J_{NN} - D_N q - \lambda \end{pmatrix} \begin{pmatrix} u_1 \\ u_2 \\ \vdots \\ u_N \end{pmatrix} = 0.$$

For all minimal architectures, there is only one component that receives more than one input (excluding negative self-interactions). For all remaining components, the rows in the matrix will consist of only two nonzero terms: the degradation term on the diagonal, $J_{ii}$ which is therefore negative, and an additional input term from another component $J_{ij}$, whose sign depends on the nature of the input. Therefore, each of these rows will take the form

$$(J_{ii} - D_i q^2 - \lambda) u_i + J_{ij} u_j = 0.$$

By definition, for a DDI $\lambda$>0 and meaning that $J_{ii} - D_i q^2 - \lambda < 0$. Consequently, $u_i$ and $u_j$ will have the same sign, and therefore be in-phase with each other if $J_{ij}$ is positive, while they will have opposing signs and be out-of-phase for negative $J_{ij}$. Therefore, the relative phase of successive components in the positive and negative feedback loops can be inferred from the signs of interaction between them, with the exception of the interaction from a component in the negative feedback loop feeding into the positive feedback loop. Moreover, two components will be in-phase if the weight of the path between them is positive, and out-of-phase if negative, unless the path passes from the negative feedback loop into the positive feedback loop, in which case the relationship will be reversed.

#### *4.2: Responses to perturbation of minimal topologies*

Having established minimal conditions for DDI, we next asked how these minimal systems respond to the inhibition of the different components. In particular, as these systems consist only of a single positive and a single negative feedback loop, we considered how the response to inhibition depends on the placement of a component relative to these two loops.

##### 4.2.1: Inhibition of response

For the inhibition of response to $u_1$, in supplementary note 3.1.1, conditions 3.3 and 3.4 it was shown that the response is dependent on the sign of

$$\frac{\sum_{\kappa_n \subseteq S_{\kappa_n}} (-1)^n w(\kappa_n) \det[\boldsymbol{C}(\tilde{\kappa}_n)]}{\det[\boldsymbol{A}]}, \quad (4.2)$$

where each summand corresponds to a different feedback loop through $u_1$. For minimal topologies, the sign of each summand dependents on two features: whether the feedback loop supported by $u_1$ is a positive or negative feedback loop (given by $w(\kappa_n)$), and whether the nodes excluded from this loop support a positive feedback loop (given by $\det[\boldsymbol{C}(\tilde{\kappa}_n)]$). For each summand, $w(\kappa_n)$ will be negative if it represents a negative feedback loop and positive for a positive feedback loop. However, the relationship between the sign of $\det[\boldsymbol{C}(\tilde{\kappa}_n)]$, and the feedback loops contained depends on the number of nodes in the system described by submatrix $\boldsymbol{C}(\tilde{\kappa}_n)$ (see supplementary note 1). Considering all possible combinations of odd and even numbers of components in matrix $\boldsymbol{A}$ and submatrix $\boldsymbol{C}(\tilde{\kappa}_n)$, it can be seen that $(-1)^n \det[\boldsymbol{C}(\tilde{\kappa}_n)] / \det[\boldsymbol{A}]$ will always be negative if submatrix $\boldsymbol{C}(\tilde{\kappa}_n)$ contains a single positive feedback loop, and will be positive if it only contains negative feedback loops.

As minimal systems only contain a single positive feedback loop and a single negative feedback loop, submatrix $\boldsymbol{C}(\tilde{\kappa}_n)$ will always contain only negative feedback loops as the two loops must share at least one component. Therefore, each summand in 4.2 will be positive if it represents a positive feedback loop and negative if it represents a negative feedback loop. For certain placements in a minimal network, $u_1$ will only fall in one feedback loop, and there will be a single nonzero summand in equation 4.2. Therefore, if $u_1$ is only found in the positive feedback loop it will decrease on its inhibition, and if it is found only in the negative feedback loop it will increase on its inhibition. However, for some placements in a minimal network $u_1$ will be in both positive and negative feedback loops. In these instances, there will be two summands in 4.2, one positive and the other negative. The response of the system will depend on the relative magnitude of each term, with both an increase and a decrease in $u_1$ possible. These constraints are summarized in fig 4b.

4.2.2: Inhibition of production

As shown in supplementary note 3.1.2, conditions 3.5 and 3.6, the response of $u_1$ to the inhibition of its own production is determined by the sign of $\det[\boldsymbol{C}(\tilde{\kappa}_1)]$ relative to $\det[\boldsymbol{A}]$. This in turn, depends on whether the system described by the submatrix excluding $u_1$ contains the positive feedback loop of the minimal network or not. If $u_1$ is excluded from the positive feedback loop, the submatrix determinant $\det[\boldsymbol{C}(\tilde{\kappa}_1)]$ will therefore contain the single positive feedback loop in the system. As matrix $\boldsymbol{C}(\tilde{\kappa}_1)$ by definition has one fewer component than matrix $\boldsymbol{A}$, as detailed in equation 3.1, $\det[\boldsymbol{A}]$ and $\det[\boldsymbol{C}(\tilde{\kappa}_1)]$ will have the same sign. However, if $u_1$ is included from the positive feedback loop, regardless of whether it is also part of the negative feedback loop, $\det[\boldsymbol{C}(\tilde{\kappa}_1)]$ will only represents negative feedback loops, and $\det[\boldsymbol{A}]$ and $\det[\boldsymbol{C}(\tilde{\kappa}_1)]$ will have opposite signs. Therefore, if $u_1$ is excluded from the positive feedback loop, it will increase upon inhibition its production, while if it is part of the positive feedback loop (whether or not it is also in the negative feedback loop) it will decrease if its production is inhibited. These constraints are summarized in fig 4b.

4.2.3: Response of $u_2$

Conditions for the response of component $u_2$ to inhibition of component $u_1$ were defined in supplemental note 3, conditions 3.7 and 3.8. Writing these in terms of paths between $u_1$ and $u_2$ (see equation 3.9) the response of $u_2$ depends on the sign of

$$-\sum_{p_n \subseteq S_{p_n}} (-1)^n w(p_n) \det[\boldsymbol{C}(\tilde{\kappa}_n)] \sum_{j=1}^{N} \mathrm{C}[\boldsymbol{A}]_{j1} b_j, \quad (4.3)$$

where each summand contains a different path from $u_1$ to $u_2$. Written in this form, it is apparent that the response of $u_2$ to the inhibition of $u_1$ is again dependent on two topological features of the minimal system: the weight of the path from component $u_1$ to $u_2$ (given by the term $w(p_n)$) and whether the nodes excluded from this path support a positive feedback loop (given by $\det[\boldsymbol{C}(\tilde{\kappa}_n)]$).

If there only exists a single path from $u_1$ to $u_2$, there will only be a single nonzero term in 4.3. As shown in the above analysis of the phase relationship (supplementary note 4.1.2), the sign of the path $w(p_k)$ will be positive if $u_1$ to $u_2$ are in-phase and negative if out-of-phase. The only exception is when $u_1$ is excluded from the positive feedback loop but the path to $u_2$ passes into or through the positive feedback loop, in which case the relationship between this relationship is reversed. As detailed in equation 3.1 the relationship between the sign of $\det[\boldsymbol{C}(\tilde{\kappa}_n)]$, and the feedback loops contained depends on the number of nodes in the system described by submatrix $\boldsymbol{C}(\tilde{\kappa}_n)$ (see supplementary note 1). Again, considering all possible combinations of odd and even numbers of components in submatrices $\boldsymbol{C}(\tilde{\kappa}_n)$ and path $w(p_k)$, and given that $\sum_{j=1}^{N} \mathrm{C}[\boldsymbol{A}]_{j1} b_j$ has the same sign as $\det[\boldsymbol{A}]$ (see supplementary note 2) the term $-(\sum_{j=1}^{N} \mathrm{C}[\boldsymbol{A}]_{j1} b_j)(-1)^n \det[\boldsymbol{C}(\tilde{\kappa}_n)]$ will be positive if $\det[\boldsymbol{C}(\tilde{\kappa}_n)]$ contains the destabilising positive feedback loop and negative otherwise.

Therefore, considering all possible placements of $u_1$ and $u_2$ relative to the two loops of the minimal system it can be seen that if $u_1$ is in the positive feedback loop (and therefore the positive feedback loop is not found in submatrix $\boldsymbol{C}(\tilde{\kappa}_n)$), $u_2$ will decrease if it is in-phase with $u_1$, and increase if out-of-phase. However, if $u_1$ is in excluded from the positive feedback loop, $u_2$ will increase if it is in-phase with $u_1$, and decrease if out-of-phase as either the path to $u_2$ will pass from the negative feedback loop into the positive feedback loop, or this path will exclude components from the positive feedback loop. These constraints are summarized in fig 4b.

It should be noted that in certain instances when $u_1$ and $u_2$ lie in both the positive and negative feedback loop, two paths can be traced from $u_1$ to $u_2$: one consisting of interactions only found in the positive feedback loop, and the other comprising only found in the negative feedback loop. There will therefore be two summands in equation 4.3, one associated with each path, and they will be of opposing signs. Therefore, the response of $u_2$ to the inhibition of $u_1$ will depends on the relative weight of each path, and will either increase or decrease.

It should be noted that while the response of $u_2$ to the inhibition of $u_1$ is not constrained by topology in these instances, it is coupled with the responses of other components in the system. Factoring out all coefficients shared between the two summands leaves

$$\left(-1\right)^{w-\nu(w^+)} w\left(p_{w^+}\right) \det[\boldsymbol{W}(\tilde{\kappa}_{w^+})] + \left(-1\right)^{w-\nu(w^-)} w\left(p_{w^-}\right) \det[\boldsymbol{W}(\tilde{\kappa}_{w^-})], \quad (4.4)$$

where matrix $\boldsymbol{W}$ is the submatrix $\boldsymbol{A}(\gamma_w)$ comprising the $w$ components supporting the two paths defined by $\gamma_w$. The terms $w(p_{w^+})$ and $w(p_{w^-})$ give the weights of the paths through the positive and negative loops respectively, $\nu(w^+)$ and $\nu(w^-)$ the numbers of components in the respective paths, and $\boldsymbol{W}(\tilde{\kappa}_{w^+})$ and $\boldsymbol{W}(\tilde{\kappa}_{w^+})$ the submatrices of $\boldsymbol{W}$ comprising the components excluded from the respective paths. This same sum is also left when considering the response of $u_1$ to its own inhibition if it falls in both feedback loops, and shared terms are factored out of equation 4.2. In both instances, if the weight through the positive feedback loop is greater than the negative feedback loop, that is

$$\left|\left(-1\right)^{w-\nu(w^+)} w\left(p_{w^+}\right) \det[\boldsymbol{W}(\tilde{\kappa}_{w^+})]\right| > \left|\left(-1\right)^{w-\nu(w^-)} w\left(p_{w^-}\right) \det[\boldsymbol{W}(\tilde{\kappa}_{w^-})]\right|$$

then $u_1$ will decrease on its own inhibition, as will any in-phase components, with out-of-phase components increasing. However, when the inequality is reversed and $u_1$ increases on its own inhibition, the response of any component to which the two paths can be traced will be coupled, increasing if it is in-phase, and decreasing if out-of-phase.

***4.3: Adding nodes to minimal topologies***

In the above analysis of the criteria for a minimal topology (supplementary note 4.1.1) we only considered strongly-connected networks, where the two feedback loops comprise all $N$ components. For systems where the two loops only pass through a subset of components, if the remaining components respond passively (i.e. do not feed back into the system) each component will only add a single interaction. Like the strongly-connected networks these networks also have two loops and $N$+1 interactions.

These passive components can be wired back into the minimal RD 'core' by adding interaction from the passive components back to the core. If the weights of any additional interactions, or the weights of additional loops generated by them, are not excessively large, the criteria for DDI will still be met, and the phase relationship of the related not strongly-connected minimal topology will also be maintained. While the resulting topologies are not strictly minimal in terms of the number of loops or interactions, they comprise additional strongly-connected networks which satisfy the conditions for DDI. Moreover, as the removal of an interaction will either result in the system either no longer satisfying the criteria for DDI or non-longer being strongly-connected, we will also consider such topologies as an additional form of minimal topology.

We will now consider the effect of inhibition on networks with additional components wired in outside of the RD 'core'. We first will consider the case where there is a single component $u_1$, external to the core and the effect of inhibiting this component on its own levels, before considering the effect of inhibiting $u_1$ on core components. We then consider the effect of external interactions on the core.

4.3.1: Effect on self

The addition of a single component external to the RD core will generally result in the addition of a single positive feedback loop to the system (we will discuss below the subset of cases where the addition of a single component results in the addition of two loops).

*Inhibition of response*

As for the minimal topologies (supplementary note 4.2.1) when the response to $u_1$ is inhibited, the behaviour of $u_1$ depends on the sign of the sum in 4.2. This sum will consist of a single term, and as before, the sign of this term depends in part on whether or not submatrix $\boldsymbol{C}(\tilde{\kappa}_n)$ contains the destabilising core positive feedback loop, and also whether this loop supports positive or negative feedback. We will ask how the wiring of this loop relative to the core positive feedback loop, along with the weighting of the loop, determines the response to inhibition.

We will first consider when the feedback loop through $u_1$ includes components from the core positive feedback loop. In this case submatrix $\boldsymbol{C}(\tilde{\kappa}_n)$ contains only negative feedback loops and $(-1)^n \det[\boldsymbol{C}(\tilde{\kappa}_n)] / \det[\boldsymbol{A}]$ will be positive. If $u_1$ itself supports a positive feedback loop condition 3.4 will be satisfied and $u_1$ will decrease on its own inhibition. Conversely, if $u_1$ supports a negative feedback loop, condition 3.3 will be satisfied and $u_1$ will increase on its own inhibition. These relationships are reverse if the loop through $u_1$ does not pass through any components in the core positive feedback loop. If this is the case, submatrix $\boldsymbol{C}(\tilde{\kappa}_n)$ will contain the core positive feedback loops and $(-1)^n \det[\boldsymbol{C}(\tilde{\kappa}_n)] / \det[\boldsymbol{A}]$ will be negative (otherwise conditions for DDI cannot be met – see supplemental note 1). In this instance, if $u_1$ supports a positive feedback loop, condition 3.3 will now be satisfied and $u_1$ will increase on its own inhibition while if $u_1$ supports a negative feedback loop condition 3.4 will be satisfied and $u_1$ will decrease on its own inhibition.

Interestingly, the response of $u_1$ to its inhibition does not correlate directs with the feedback loop $u_1$ supports, but rather with the net feedback provided to the core positive feedback loop (ie the weight the path from $u_1$ to a core positive feedback components multiplied by the weight of the return path). If $u_1$ supports a net positive feedback, it will decrease on its own inhibition, while if it provides a net negative feedback, it will increase on its own inhibition. These constraints are summarized in fig 4d.

*Inhibition of production*

As demonstrated in supplementary note 3, the response to the inhibition of production is dependent on the sign of det $[\boldsymbol{C}(\tilde{\kappa}_1)]$ relative to det $[\boldsymbol{A}]$ (condition 3.5 or 3.6). As det$[\boldsymbol{A}]$ and det$[\boldsymbol{C}]$, only differ by the presence of the degradation coefficient $c_1$ in the first matrix term it follows that

$$1 = \frac{-(-1)^n w(\kappa_n)\det[\boldsymbol{C}(\tilde{\kappa}_n)] - c_1 \det[\boldsymbol{C}(\tilde{\kappa}_1)]}{\det[\boldsymbol{A}]}.$$

In the above analysis of the inhibition of response, we demonstrated how network topology (i.e. the sign of the loop containing $u_1$, and how this feeds into the core RD network) determines the sign of $(-1)^n w(\kappa_n)\det[\boldsymbol{C}(\tilde{\kappa}_n)]/\det[\boldsymbol{A}]$. From these arguments, it follows that if $u_1$ supports a net positive feedback, $-c_1\det[\boldsymbol{C}(\tilde{\kappa}_1)]/\det[\boldsymbol{A}]$ must be negative. As $-c_1$ is negative, condition 3.6 will be satisfied and $u_1$ will decrease. However, if $u_1$ supports a net negative feedback, the sign of $-c_1\det[\boldsymbol{C}(\tilde{\kappa}_1)]/\det[\boldsymbol{A}]$ is not constrained and therefore both conditions 3.5 and 3.5 can be satisfied. Therefore, $u_1$ will decrease on the inhibition of its production if it provides a net positive feedback to the core positive feedback loop, but depending on the strength of the different feedback loops, $u_1$ will either increase or decrease when its production is inhibited if it provides a net positive feedback. These constraints are summarized in fig 4d.

It should be noted that in some instances where $u_1$ feeds into and out of the core RD network at components which are found in both core positive and negative feedback loops, it is possible that a second loop supported by $u_1$, through components only found in the core negative feedback loop, will also exist. In this case the two terms describing alternate paths through the positive feedback loop and negative loop (see 4.4) will again remain after shared components are factored out. As discussed above, in the cases where the magnitude of the path through the negative feedback loop is greater than the path through the positive feedback loop, the relationships between the response of $u_1$ to its own inhibition and the feedbacks it supports described here will also be inverted.

4.3.2: Effect on core components

We will now consider the how inhibiting the external component $u_1$ affects a core component $u_2$. We will consider the effect of whether $u_1$ feeds directly into the positive or negative feedback loop of the core RD network, and on whether $u_2$ itself lies in the core positive or negative feedback loop. As for minimal systems (supplementary note 4.2.2), $u_2$ will decrease on the inhibition of $u_1$ if the sign of the path from $u_1$ to $u_2$ is positive, unless the submatrix determinant det $[\boldsymbol{C}(\tilde{\kappa}_n)]$ contains the positive feedback loop of the core RD network. Likewise, $u_2$ will increase if the path it is negative, unless the path excludes the core RD network positive feedback loop.

If $u_1$ feeds into the core RD network through the positive feedback loop, det $[\boldsymbol{C}(\tilde{\kappa}_n)]$ will not contain the core positive feedback loop. The response will therefore depend on the sign of $w(p_k)$, such that if $u_1$ itself supports a positive feedback loop. The weight of the path from $u_1$ to $u_2$ will be positive if $u_1$ and $u_2$ are in-phase, and negative if they are out-of-phase. Therefore, $u_2$ will decrease if it is in-phase with $u_1$ and increase if it is out of phase. The opposite will occur if $u_1$ is in a negative feedback loop, with the level of $u_2$ increasing if it is in-phase with $u_1$ and decreasing if it is out-of-phase.

We next consider when $u_1$ feeds back into the core through the negative feedback loop. If $u_2$ is also found in this loop, and the path from $u_1$ does not pass through the core positive feedback, the sign of the path will be positive if the components are in-phase and negative if they are out-of-phase provided $u_1$ supports a positive feedback loop (and therefore provides a net negative feedback to the core positive feedback loop). The reverse is true if $u_1$ supports a negative feedback loop (and thus a net positive feedback). As the core positive feedback components are found in det $[\boldsymbol{C}(\tilde{\kappa}_n)]$, if $u_2$ is in-phase with $u_1$ it will decrease on inhibition if $u_1$ provides a net positive feedback, and out-of-phase components will increase, with the reverse true if $u_1$ provides a net negative feedback. The same is also true when $u_1$ feeds into the core negative feedback loop, but the path to $u_2$ passes into or through the core positive feedback loop. When the core positive feedback components are found no longer in submatrix $\boldsymbol{C}(\tilde{\kappa}_n)$ and therefore the sign of det $[\boldsymbol{C}(\tilde{\kappa}_n)]$ is reversed, the relationship between the signs of the path from $u_1$ to $u_2$ and the phase relationship are now inverted by passing back into the core positive feedback loop, maintaining the above relationship.

Therefore, regardless of where the path from $u_1$ feeds into the core RD network, if $u_1$ net negative feedback loop, $u_2$ will increase if it is in-phase with $u_1$, and decrease if it is out-of-phase. The opposite will be true if $u_2$ supports a net positive feedback. These constraints are summarized in fig 4d. Again, there can be instances where two paths of opposing sign can be traced from $u_1$ to $u_2$ if $u_1$ feeds into a component found in both core loops and $u_2$ is also found in both loops. The response is again dependent on the sign of equation 4.4, with the above responses inverted if the magnitude of the term relating to the path through the negative feedback loop is greater than those relating to the positive feedback loop, with all components receiving input from the two paths responding in a coupled manner.

4.3.3: Effect of adding external components to RD system

Finally, we considered the effect adding external components to existing constraints on the system. We asked whether the addition of an external component could change the behaviours within the minimal RD core. As shown above, a major feature determining the response of a component in a minimal RD network either to its own inhibition, or the inhibition of another component, is the whether or not various submatrices contain the destabilizing positive feedback loop of the core RD network. In some instances, the addition of loops can change the stability of these submatrices, either making stable submatrices unstable by adding a positive feedback loop, or making unstable submatrices stable by the addition of further negative feedback. However, it should be noted that as the various loops and paths that determine the behaviours within the core will still be present, only certain placements of feedback loops have such effects, and that they only affect certain components under particular parameterisations. There will still exists parameterisations where the responses described above (summarized in fig 4b) are maintained.

The constraints applying to the addition of one external component will, therefore, apply to any number of external components, provided that the magnitudes of additional interactions are not sufficiently large so as to change the stability of any submatrices. This also opens the possibility of considering potential interactions between external components $u_1$ and $u_2$ (for simplicity we will only consider the cases where the path between $u_1$ and $u_2$ passes through the core). The path will contain the path from $u_1$ to a core component, plus an additional interaction to $u_2$. Therefore, the response of $u_2$ to the inhibition of $u_1$ will be the same the preceding core component in the path if the final interaction is positive, and the opposite if it is negative. The same is true of the response of external $u_2$ to a core $u_1$. Consequently, if $u_1$ is involved in positive feedback (either as the core loop or as a source of additional positive feedback), then $u_2$ will decrease if it is in-phase with $u_1$, and increase if it is out-of-phase, while if $u_1$ provides negative feedback the reverse is true. As has been discussed above, when two paths can be traced from $u_2$ to $u_1$, if the path through the negative feedback loop has a greater weighting these relationships will be inverted coupled to other such components.

Taken together, the effects of inhibiting $u_1$ on itself, and on $u_2$ primarily depend on whether $u_1$ falls in the core positive feedback loop, or provides an additional net positive feedback, or whether it provides negative feedback. While for some parameterisations of some topologies, certain components will behave differently, these constraints on the behavior of minimal systems provide a framework to consider the behavior of RD networks in response to inhibition experiments.


Desoer, C. A. (1960). "The Optimum Formula for the Gain of a Flow Graph or a Simple Derivation of Coates' Formula." Proceedings of the IRE **48**(5): 883-889.

Marcon, L., X. Diego, J. Sharpe and P. Muller (2016). "High-throughput mathematical analysis identifies Turing networks for patterning with equally diffusing signals." Elife **5**.

Murray, J. D. (2003). Mathematical Biology, Berlin: Springer-Verlag.

White, K. A. J. and C. A. Gilligan (1998). "Spatial heterogeneity in three species, plant–parasite–hyperparasite, systems." Philosophical Transactions of the Royal Society of London. Series B: Biological Sciences **353**(1368): 543-557.
